# Anthropogenic food enhancement alters the timing of maturational landmarks among wild savanna monkeys (*Chlorocebus pygerythrus*)

**DOI:** 10.1101/2020.04.18.048314

**Authors:** Christopher A. Schmitt, Alicia M. Rich, Stacy-Anne R. Parke, Maryjka B. Blaszczyk, Jennifer Danzy Cramer, Nelson B. Freimer, J. Paul Grobler, Trudy R. Turner

## Abstract

Anthropogenic landscapes are rapidly replacing natural nonhuman primate habitats. Yet, the access to anthropogenic resources on primate biology, health, and fitness remain poorly studied. Given their ubiquity across a range of human impacted landscapes, from cities to national parks, savanna monkeys (*Chlorocebus* spp.) provide an excellent study system in which to test these effects. We compared body condition and reproductive maturation in vervets (*Chlorocebus pygerythrus*) inhabiting a private farm in !Gariep Dam, with ample access to anthropogenic foods, and wild-foraging vervets in Soetdoring Nature Reserve, South Africa. Overall, vervets in !Gariep show significantly thicker skin folds, and higher BMI and body mass, than those in Soetdoring, suggesting increased fat deposition. Males in !Gariep have larger relative testis volumes at peri-pubescent ages compared to those in Soetdoring, suggesting early reproductive maturation associated with age-specific increases in body mass. Females from !Gariep showed evidence of an earlier onset of reproduction than those in Soetdoring, based on parity status as assessed by nipple length and evidence of lactation. Parity status at sub-adult dental ages was also strongly associated with body mass. These results are consistent with a positive effect of anthropogenic food-enhancement on body fat deposition, potentially linked to an earlier onset of reproductive maturation. Further investigation into primate responses to cultivated resources will inform our understanding of the broader effects of food enhancement on developmental plasticity.

## INTRODUCTION

Research on primate life histories often focuses on inter-specific variation across the order. In these studies, variation presumably reflects adaptive responses to long-sustained predation risks, dietary constraints, demographic variables, or other long-term socioecological factors (Hill 2005; Kamilar & Cooper 2013). Within species, however, life histories may also be highly plastic in response to the energetic constraints of the uterine and postnatal environment. Testicular tissue generation and spermatogenesis are energetically costly developmental processes (Rato et al. 2012), and female reproduction has been well-established as energy-limited (Pusey 2012). The initiation of puberty in both males and females, for example, is controlled indirectly by nutrition and adiposity via the permissive effects of the adipose-derived hormone leptin (Elias 2012). Undernutrition *in utero* delays testis growth and puberty in male offspring (Zambrano et al. 2014). While maternal undernutrition delays ovarian development and menarche in females, maternal overnutrition leads to early menarche (Zambrano et al. 2014). Overall, increased nutrient availability during postnatal development accelerates the onset of reproductive traits, including ovarian function and menarche, testis development, the emergence of secondary sexual characteristics, and subsequent onset of reproduction (Ellison 1990; Koziel and Jankowska 2002; Setchell & Lee 2004; Gluckman and Hanson 2006a).

Given this, it is not surprising that primate populations living in nutrient- or calorie-rich environments consistently show relatively rapid life histories for their species, including earlier ages at sexual maturity and first reproduction (Altmann & Alberts 2003; Kuzawa & Bragg 2012). With the rapid expansion of human landscapes (Estrada et al. 2017), crop foraging and provisioning are growing sources of food for wild non-human primates (Naughton-Treves et al. 1998; Strum 2010; Lodge et al. 2013; Hill 2017). These anthropogenic foods are often more accessible than wild forage (Altmann & Muruthi 1988; Cancelliere et al. 2018), allowing primates to save energy that they might otherwise spend searching for and processing food. Crops may also be higher in available energy content (Saj et al. 1999, 2001; Riley et al. 2013), allowing for an earlier threshold of reproductive maturation. While ecologically-mediated malnutrition can slow reproductive timing and introduce life-long constraints on reproduction (Bercovitch & Strum 1993; Lea et al. 2015), anthropogenic foods may provide a release from such constraints. Human-provisioned yellow baboons, for example, grow one-third faster and to almost twice the size of their wild-foraging counterparts (Altmann & Alberts 2005; Onyango et al. 2013). Similarly, crop foraging baboons in Gashaka-Gumti National Park in Nigeria showed shorter inter-birth intervals and lower infant mortality (Higham et al. 2009), while Japanese macaques (*Macaca fuscata*) also began giving birth at younger ages during periods of nutritional provisioning (Mori 1979).

Savanna monkeys (*Chlorocebus* spp.) adapt well to anthropogenically impacted habitats throughout their range (Brennan et al. 1985; Saj et al. 2001; Lee & Priston 2005). Wild populations are ubiquitous across sub-Saharan Africa and can also be found in a range of ecologies, from nature reserves to intensive agricultural areas, with varying proximity to anthropogenic landscapes (Fourie et al. 2015; Turner et al. 2018). They are also particularly well-characterized behaviorally, genomically, and physiologically, both in the wild (Jasinska et al. 2013; Svardal et al. 2017; Turner et al. 2018) and in captivity (Kavanagh et al. 2007; Schmitt et al. 2018).

The growth and reproductive ecology of savanna monkeys has also been well-characterized. Wild savanna monkeys show a menstrual cycle lasting a median of 33 days (range: 25-46 days; *Ch. pygerythrus* near Kampala, Uganda; Rowell 1970). Although females may mate throughout their cycle (Rowell 1970; Andelman et al. 1985), most savanna monkeys have a distinct mating season from April through July (McFarland et al. 2014; Blasczcyk 2016). As such, pregnancy and birth is largely seasonal, with most births in South African populations occurring between October and December (Cheney et al. 1988; Blaszczyk 2016, Jarrett et al. 2020). In wild vervet monkeys, lactation begins during pregnancy and lasts between 9 and 18.5 months after birth (Whitten 1982; Lee 1984). Provisioned vervets have a shorter time to weaning and earlier subsequent conceptions (Whitten 1982). Birth cohorts of wild-feeding South African vervet monkeys (*Ch. pygerythrus*) reach adult size more slowly and at a later age than those in captivity (Jarrett et al. 2020). In wild green monkeys (*Ch. sabaeus*) in St. Kitts & Nevis, this leads to clear increases in body mass and better body condition in captive adults (Turner et al. 2016). Relatively malnourished wild cohorts also grow more slowly and to a lighter adult weight than those with ample food (Jarrett et al. 2020). Wild male vervets reach pubertal landmarks beginning between 23 and 37 months with the descent of the testes, followed by detectable spermatogenesis at 48-60 months (Whitten & Turner 2009), and ejaculatory copulation at 60 months of age (Cheney et al. 1988). Dispersing males typically leave their natal group between 48 and 84 months, with the majority leaving by 72 months (Cheney & Seyfarth 1983). Females do not give birth until 52-68 months, although vervets provisioned with human food waste or living in resource rich environments tend to give birth at earlier ages (Brennan et al. 1985; Cheney et al. 1988). The ability to bring an infant to term in wild female vervets is contingent on both age and rank. Both very young mothers and older females have a higher probability of miscarriage (Turner et al. 1987). Low ranking females, with presumably lower priority of access to resources, are unable to reproduce annually (and the lowest ranked for multiple years) unlike those of higher rank (Turner et al. 1987). In all previous comparative studies among vervet populations, stark differences in attaining growth and maturational landmarks were noted based on resource availability and quality (Cheney et al. 1988; Turner et al. 1997; Whitten & Turner 2009; Turner et al. 2018; Jarrett et al. 2020).

Here we compare aspects of reproductive maturation and body condition in two populations of vervet monkeys in South Africa, each living in similar biomes with contrasting human impacts. The first population, in Soetdoring Nature Reserve, subsists primarily on natural forage while experiencing only moderate human impacts (Blasczyzk 2016). The second population, on private farms near the !Gariep Dam, has ample access to anthropogenic foods and experiences high human impacts. We predicted that food-enhanced vervets living on private farms near the !Gariep Dam, compared to those living in the Soetdoring Nature Reserve, would exhibit (1) body condition consistent with increased nutritional enhancement, including heavier body weight, higher body mass index (BMI), and thicker skin folds, and (2) associated evidence of earlier reproductive maturation, as illustrated by (2a) an earlier increase in testis volume and pubertal growth spurt in males, and (2b) earlier average age at first birth in females, using nipple morphology as an indicator of parity.

## METHODS

### Study Sites

Both Soetdoring Nature Reserve and the !Gariep Dam farms occupy near-identical grassland biomes (Janecke 2002; Janecke & du Preez 2005), at similar altitudes (1261 masl and 1206 masl), and with similar levels of annual precipitation (79 mm and 72 mm) and temperature (16.7° C and 16.9° C) over the past decade (Fig 1a). In both locations, vervet groups primarily occupy riparian forest and adjacent *Acacia* thornveld (Blaszczyk 2016), although the vervets in !Gariep spend much of their time in agricultural fields and orchards.

**Figure 1.**
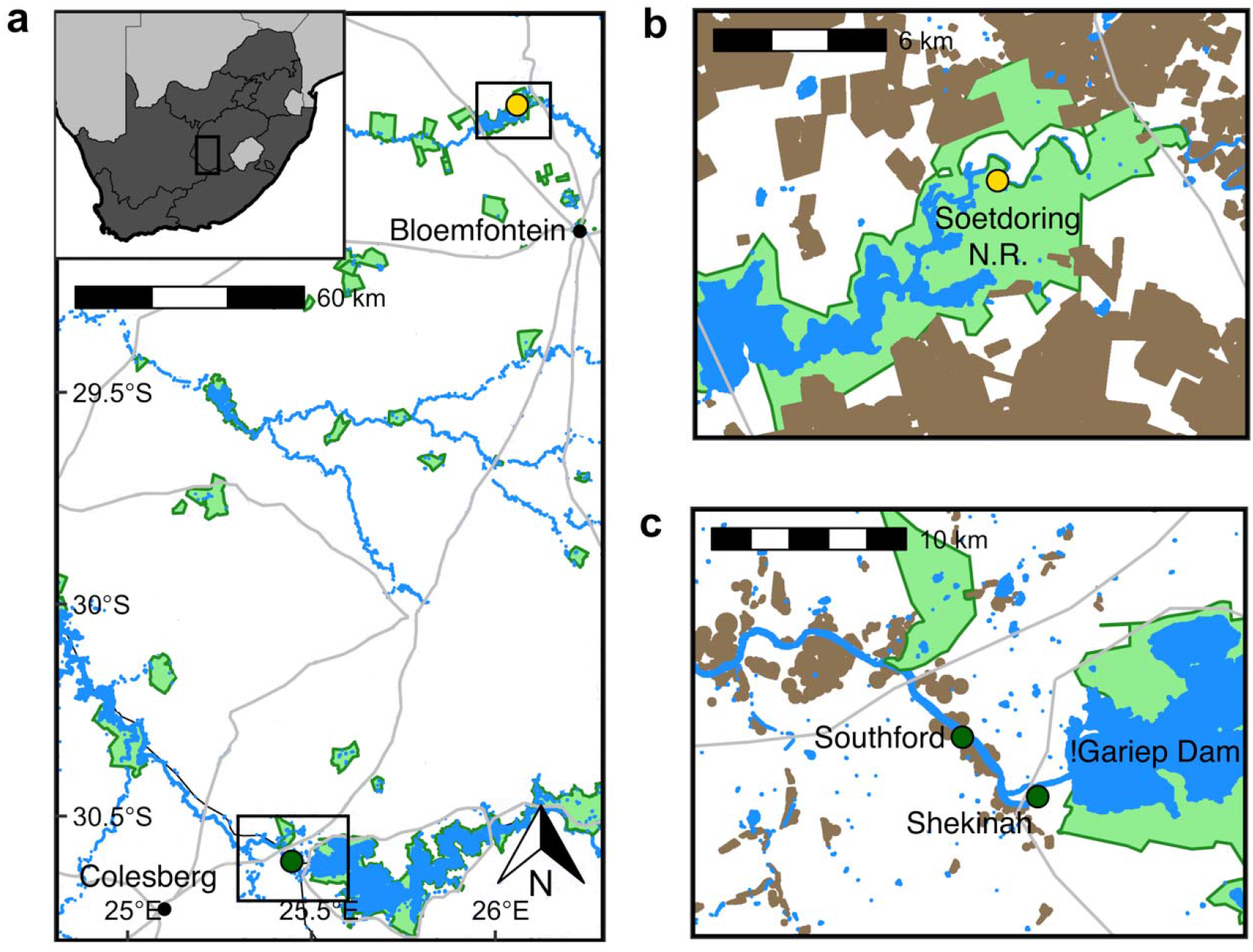
Study sites in South Africa (a), with insets of b) Soetdoring Nature Reserve and c) the !Gariep Dam region. Brown areas indicate cropland, blue include rivers, and both naturally and dammed water sources, while green are protected natural areas. Soetdoring Nature Reserve (gold circle) has fewer human-wildlife interactions and provisioning, while private farms near the !Gariep Dam (green circles) facilitate crop foraging and high calorie provisioning. Both study areas include mosaics of grassland and Nama Karoo and have near identical ecological conditions outside immediate human impacts within their ranges.

*Soetdoring Nature Reserve* (Fig 1b) occupies 6173 Ha of highveld grassland, including the Modder River and the Krugersdrift Dam (Janecke & du Preez 2005). The vervet groups studied here remain on the south side of the Modder River, primarily occupying riparian forest and adjacent *Vachellia* thornveld near the eastern entrance to the reserve. Although they almost exclusively subsist on natural forage, food scraps left by visitors were also eaten (Blaszczyk 2016). The landscape surrounding the reserve consists almost entirely of large-scale agricultural fields consisting of maize, wheat, alfalfa, and sunflower cultivation.

*Southford Stud* and the *Shekinah Guest Farm* are private farms near the !Gariep Dam (Fig 1c). Southford Stud is a 10,000-acre private farm primarily dedicated to horse husbandry, straddling the Free State and Northern Cape provinces along the Orange River. It contains several hundred acres of mixed pasture, riparian forest, and cropland including wheat, maize, alfalfa, and pecans. Vervets at Southford Stud are also commonly seen eating the feed provided *ad libitum* to the horses. The Shekinah Guest Farm, a 10 minute drive from Southford Stud, is a tourist guest farm on the Orange River with riparian forest and small agricultural holdings, including wheat and maize.

### Field Collections

We trapped and sampled monkeys at both sites across four trapping seasons between May and August in 2010 and 2016-2018 (Supplementary Table 1). Previous publications provide greater detail on trapping and data collection methods (Jasinska et al. 2013; Turner et al. 2018). We baited vervets with maize into modified drop traps (Grobler & Turner 2010), where they were sedated by a veterinarian with 4 mg/kg of equal parts medetomidine/ketamine (2016-2018) or ketamine/xylazine (2010). We placed subdermal microchips in the interscapular region to facilitate identification. We collected morphological data as described by Turner et al. (1997; 2018): we used measuring tape for lower leg length (as the best proxy for body size in these data, in keeping with Rodriguez et al. 2015), body length (for BMI, measured as body mass in kg divided by body length in m squared) and waist circumference, a digital scale to measure body mass, Lange calipers for skinfold thickness (mid-biceps, supra-iliac, sub-scapular, and peri-umbilicus), and an ochidometer for testis volume (Karaman 2005; Cramer et al. 2013). We assigned dental age categories and assessed approximate chronological ages using dental eruption patterns as described in Turner et al. (2018) (Table 1). We assigned female parity status based on nipple morphology (Altmann et al. 1981) and used digital photographs of the torso taken during trapping to confirm field-based assignments. Female parity status was determined based on nipple length as defined by Turner et al. (1997): nulliparous, wherein nipples are flat to the chest; primiparous, wherein nipples are firm and noticeably protrude less than a centimeter from the body; and multiparous, wherein nipples are limp and extend over a centimeter from the body. We used transabdominal or rectal palpation to assess pregnancy status (Turner et al. 1987; Eley 1992), and assessed lactation by the presence of expressible milk from the nipple (Whitten & Turner 2009). Given that collections occurred over the first few months after the breeding season, and that the earliest potential detection of pregnancy using these methods in captive primates is ~30 days post-conception (van Pelt 1974; Eley 1992), it is possible that females in our non-pregnant sample were in the early stages of pregnancy but not detectable. We urge caution in interpreting the results specific to pregnancy status given that maternal weight gain in primates can begin before the time when pregnancy is detectable by palpation (e.g., in macaques: Kohrs et al. 1976; Kemnitz et al. 1984).

**Table 1:** Dental age categories based on tooth eruption sequences in Chlorocebus (from Turner et al., 2018). Age range listed is the lower range for the initiation of that age class.

| Dental age class | Permanent dentition present | Age range (months) |
| --- | --- | --- |
| 1 | All deciduous | 6 – 115 days |
| 2 | M1 | 12 – 14 |
| 3 | M1, I1, I2 | 22 – 27 |
| 4 | M1, I1, I2, M2 | 26 – 31 |
| 5 | M1, I1, I2, M2, P3, P4 | 32 – 40 |
| 6 | M1, I1, I2, M2, P3, P4, C | 38 – 41 |
| Adult | Eruption of M3 | > 38 |

### Ethical Note

These methods have been used with great success in these populations since 2009 (Jasinska et al. 2013; Turner et al. 2019). The processing of anesthetized animals rarely lasts longer than 20 minutes, and all animals sampled in this study were successfully released back to their social groups unharmed. Consistent with South African law, a licensed South African veterinarian applied or supervised all invasive methods. The animal care and use committees at Boston University, University of California at Los Angeles, the University of Wisconsin at Milwaukee, and the University of the Free State approved all methods. All methods are consistent with the Principles for the Ethical Treatment of Non-Human Primates by the American Society of Primatologists.

### Statistical Analysis

The sample for this study includes 245 wild vervets, including 135 males and 110 females (Table 2). We included only the initial trapping data for any vervets re-trapped within or across field seasons. We conducted all analyses using R v. 3.6.1 (R Core Team 2019). Measures indicative of body condition–including body mass, BMI, waist circumference, and skinfold thicknesses–are all significantly correlated with each other. Previous work in *Macaca* also show all of these measures scale strongly with adiposity (Colman et al. 1999). Colman et al. (1999) found waist circumference to be the strongest indicator of central adiposity in *Macaca*. In our sample, however, waist circumference showed an unacceptable level of variance across sampling periods, so we discarded it. The 2010 sample lacked skinfold thickness measures, so we used body mass and BMI as indicators of adiposity in our models.

**Table 2:**
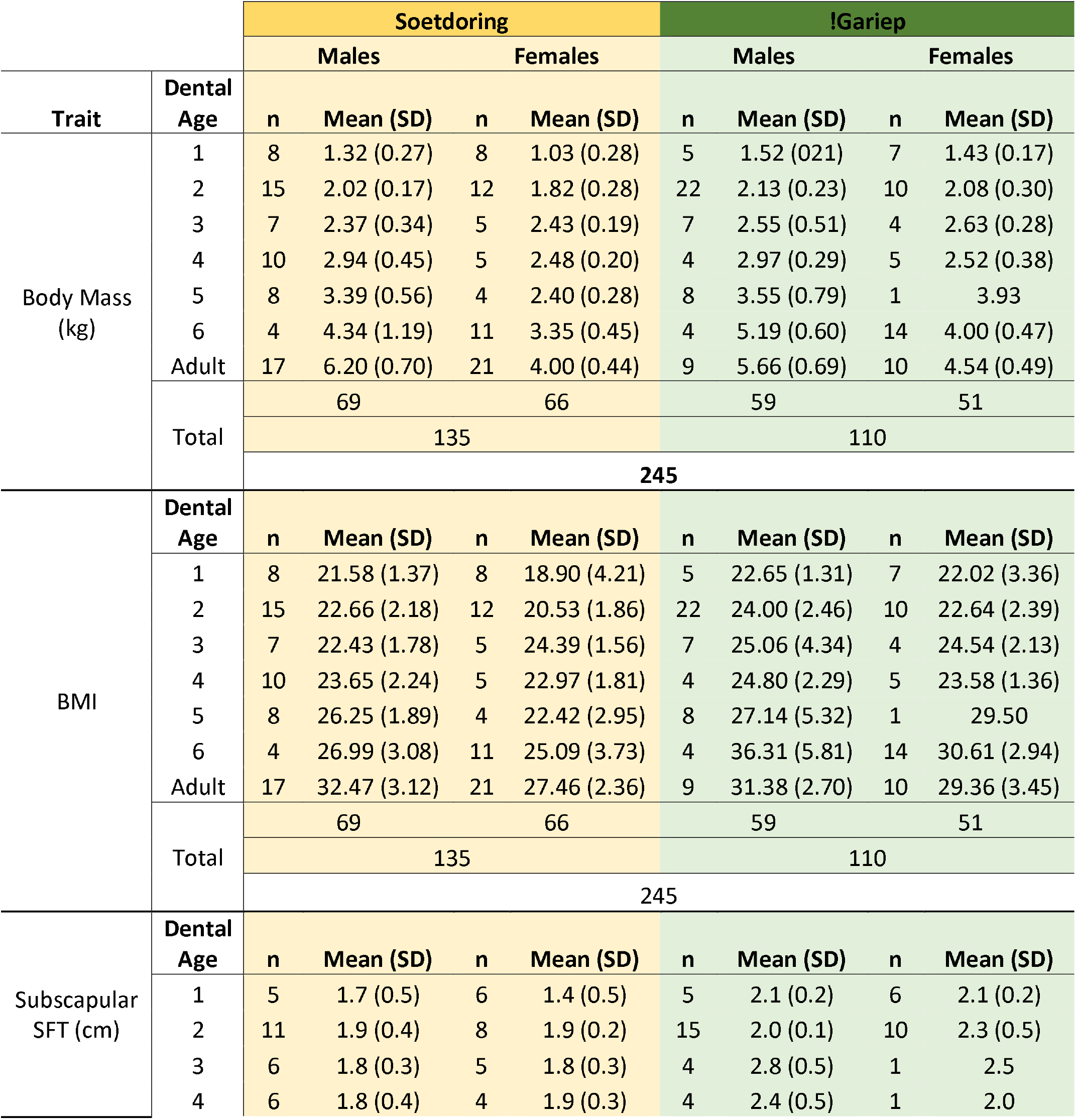
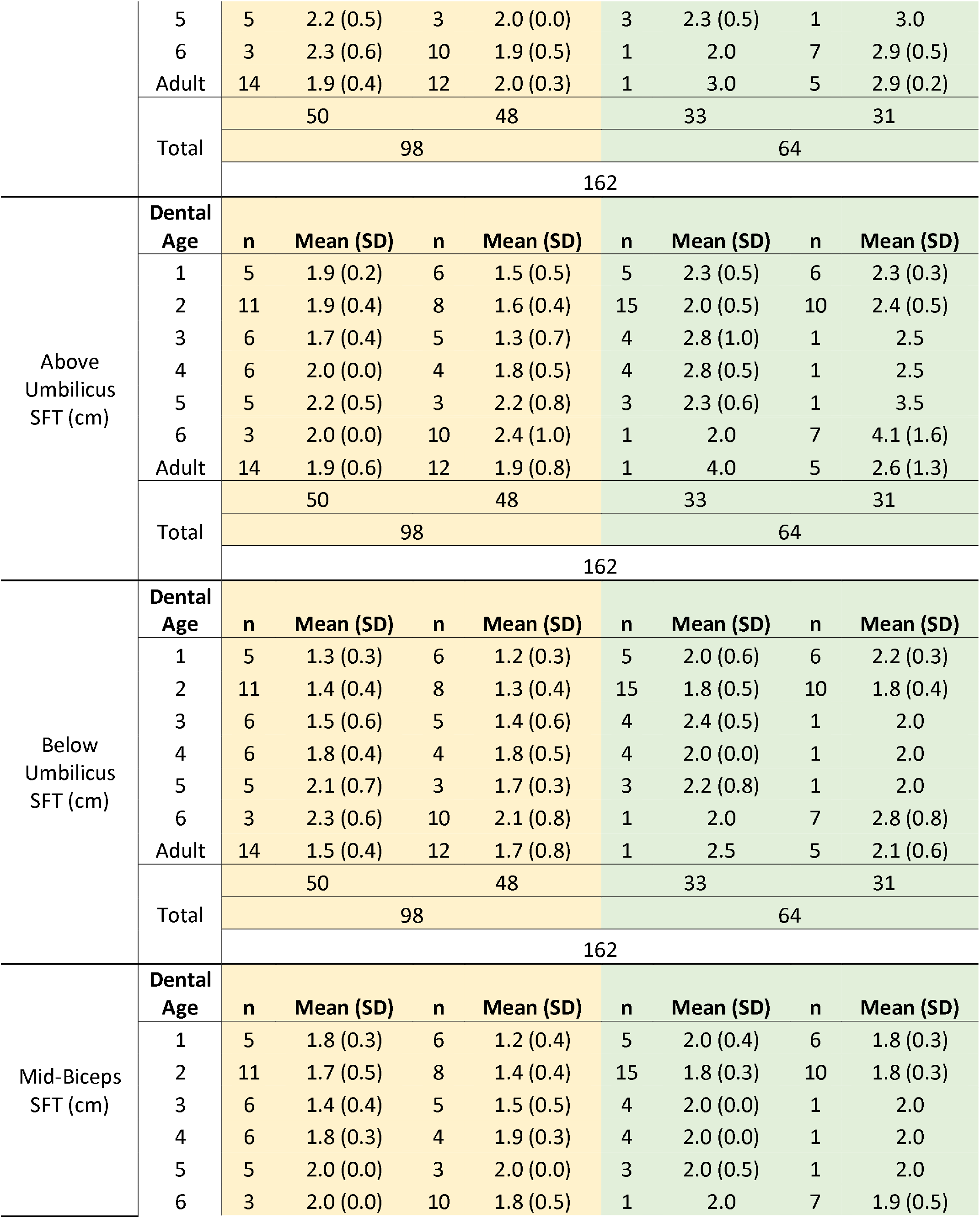
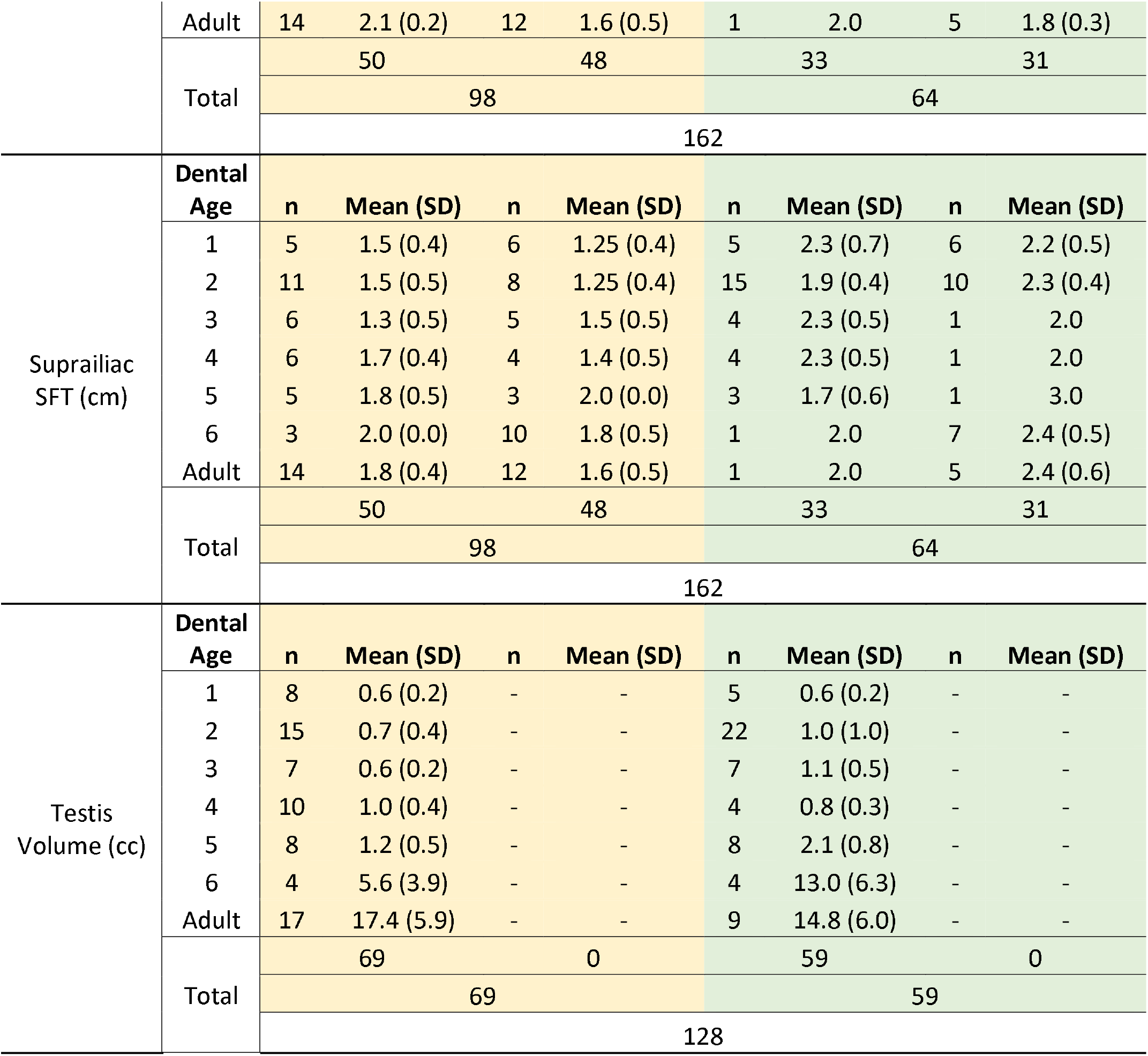
Summary statistics of vervet body measures.

We compared body measures across age, sex, and site using Welch’s ANOVA, with non-parametric Games-Howell post-hoc tests to account for differences in variance and small sample sizes across categories. We ran a separate Type II ANOVA with Games-Howell post-hoc tests on subsets of females for which pregnancy or lactation were noted in dental age 6 and adulthood to assess the impact of these statuses on body mass, BMI, and skinfold thickness. Dental age 5 was excluded from ANOVAs analyzing female variation due to low sample size in !Gariep (n = 1). Although both pregnancy and lactation status were accompanied by significantly heavier body mass and higher BMI in dental ages 6 and adulthood among these subsets (see Results and Supplemental Information), we did not include them as covariates in maturation models as doing so would have significantly reduced our sample size.

We modeled indicators of maturation in two phases: a pre-pubertal phase with relative stasis in trait state, and a peri-pubertal phase with marked change across age categories through adulthood. We placed the dividing line for these models at dental age 4 in both sexes. For males, we modeled testis volume as a continuous variable, cube root transformed to reduce the measure to one dimension, using loess curves for visualization and generalized linear regression with a Gamma error family and log link function with location, dental age, and indicators of body condition as covariates. We used the Akaike Information Criterion with a correction for small sample size (AICc) and likelihood ratio tests to assess covariate inclusion. We assessed model fit as 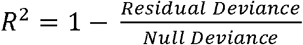. For females, there was no variation in nipple morphology prior to dental age 4, so we did not model this trait for the pre-pubertal phase. In the female peri-pubertal phase, we binned parity status into a binary variable – nulliparous vs. parous – modeled using logistic regression with location, dental age, and indicators of body condition as covariates. We also used Fisher’s exact tests to assess relative proportions of parity in each age category between sites.

### Data Availability

The final datasets generated and analyzed during this study are available from the corresponding author upon reasonable request.

## RESULTS

### Morphometric indicators of body condition

Individuals in !Gariep were, overall, significantly heavier than those in Soetdoring for both females (ANOVA: F_1,103_ = 29.68, P < 0.001), and males (F_1,114_ = 3.64, P = 0.059), although these differences only emerged beginning in the peri-pubertal phase (dental age 5, or 32-40 months of age; Fig. 2). Only one female from dental age 5 was sampled in !Gariep (body mass = 3.93 kg), making comparison across sites difficult at this age, although she weighed much more than any female sampled in that age category at Soetdoring (body mass = 2.40 ± SD 0.28 kg). At dental age 6, females in !Gariep were heavier than in Soetdoring (Games-Howell: Δ_mean_ = 0.64 kg, t = 3.50, df = 22.13, P = 0.082), and this difference continued into adulthood (Games-Howell: Δ_mean_ = 0.60 kg, t = 3.48, df = 22.91, P = 0.084). Males showed no significant differences in body mass within age categories across sites (F_6,114_ = 1.03, P = 0.407).

**Figure 2.**
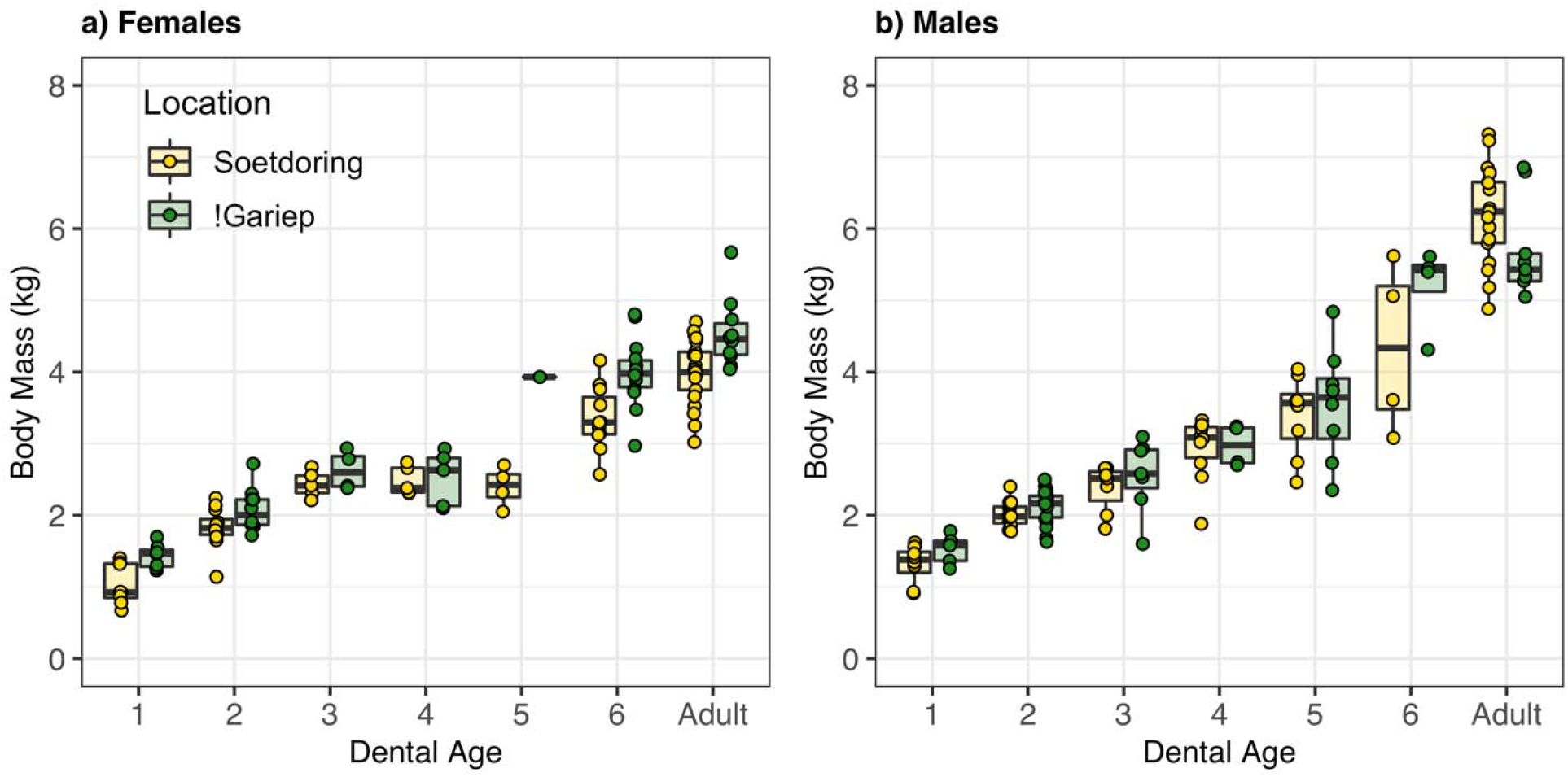
Body mass (kg) in a) female and b) male vervet monkeys across each dental age category. For estimated chronological ages, see Table 1.

Although pregnancy and lactation both were accompanied by significant increases in body mass for females (see below), site and age category showed larger effects, with adult females being significantly heavier than subadults (ANOVA: F_5,31_ = 19.67, P < 0.001; Supplementary Table 2) and females in !Gariep being significantly heavier than those in Soetdoring (F_1,31_ = 32.27, P < 0.001). Pregnant females were heavier, overall, than non-pregnant females (F_1,31_ = 5.40, P = 0.027). In dental age 6, pregnant females in Soetdoring were 7% heavier than non-pregnant females (N_preg_ = 6 of 10, Δ_mean_ = 0.25 kg), while in !Gariep pregnant females were 11% heavier than non-pregnant females (N_preg_ = 5 of 11, Δ_mean_ = 0.46 kg). Although pregnant adult females in Soetdoring were 13% heavier than non-pregnant adult females (N_preg_ = 2 of 12; Δ_mean_ = 0.54 kg), there was no detectable difference in body mass based on pregnancy in !Gariep adults (N_preg_ = 2 of 6; Δ_mean_ = 0.05 kg). Similarly, although lactating females were, overall, heavier than non-lactating females (F_1,47_ = 8.35, P = 0.006; Supplementary Table 9), these differences were small compared to those between subadults and adults (F_1,47_ = 13.22, P < 0.001) and between !Gariep and Soetdoring (F_1,47_ = 12.82, P < 0.001). There was no statistically significant difference in body mass by lactation status within each site among adults, although lactating adult females were ~6% heavier than non-lactating females in each. Lactating females in !Gariep at dental age 6 were heavier (11%) than non-lactating females (N_lact_ = 7 of 14; Δ_mean_ = 0.63 kg); no females in dental age 6 in Soetdoring were observed to be lactating.

BMI was significantly higher overall for females in !Gariep compared to Soetdoring (ANOVA: F_1,103_ = 14.47, P < 0.001), but BMI did not differ significantly between the two sites within any particular age/sex class except at dental age 6, in which !Gariep females had significantly higher BMI than those in Soetdoring (Games-Howell: Δ_mean_ = 5.52 kg/m^2^, t = 4.02, df = 18.75, P = 0.033). BMI was also significantly higher, overall, in !Gariep males than in males at Soetdoring (F_1,114_ = 5.23, P = 0.024), but there were no significant differences within age classes across sites.

Pregnant females had a significantly higher BMI, overall, than non-pregnant females (ANOVA: F_1,31_ = 4.64, P = 0.039; Supplementary Table 3), but this was overshadowed by differences between sites (F_1,31_ = 17.63, P < 0.001). Although pregnant adult females had 9% higher BMI than non-pregnant adult females in Soetdoring (Δ_mean_ = 2.7 kg/m^2^), no difference was seen in !Gariep (Δ_mean_ = 0.0 kg/m^2^). In dental age 6, pregnant females in Soetdoring had 10% higher BMI than non-pregnant (Δ_mean_ = 2.7 kg/m^2^), and in !Gariep the BMI of pregnant females was 3% higher (Δ_mean_ = 0.9 kg/m^2^). Lactating females, overall, had higher BMI than non-lactating (F_1,47_ = 4.05, P = 0.050; Supplementary Table 10). Lactating adult females in Soetdoring had 2% higher BMI (Δ_mean_ = 0.5 kg/m^2^) and in !Gariep had 13% higher BMI (Δ_mean_ = 4.1 kg/m^2^), although neither difference was statistically significant. Lactating females in dental age 6 in !Gariep had 8% higher BMI than non-lactating females (Δ_mean_ = 2.6 kg/m^2^).

All skin folds measured were significantly thicker in !Gariep compared to Soetdoring (Fig 4), except for mid-biceps in males. There were no significant age-related differences in any skin fold thickness. Overall, vervets in !Gariep showed significantly thicker skin folds, including the subscapular (males, F_1,81_ = 13.34, P < 0.001; females, F_1,78_ = 33.15, P < 0.001), above umbilicus (males, F_1,81_ = 10.63, P = 0.0016; females, F_1,78_ = 18.35, P < 0.001), below umbilicus (males, F_1,81_ = 11.10, P = 0.0013; females, F_1,78_ = 15.44, P < 0.001), mid-biceps (males, F_1,81_ = 1.34, P = 0.25; females, F_1,78_ = 8.53, P = 0.0046), and suprailiac (males, F_1,81_ = 12.97, P < 0.001; females, F_1,78_ = 50.35, P < 0.001). Pregnancy was generally not associated with thicker skin folds (Supplementary Tables 4-8, 11-15), with the notable exceptions of the mid-biceps (ANOVA: F_1,27_ = 10.00, P = 0.0039; Supplementary Table 7) and suprailiac (F_1,27_ = 4.12, P = 0.052; Supplementary Table 8), which were thicker during pregnancy in Soetdoring but thinner during pregnancy in !Gariep (F_1,27_ = 7.51, P = 0.011). The suprailiac skin fold was also thicker in Soetdoring but thinner in !Gariep in lactating compared to non-lactating females (F_1,27_ = 4.39, P = 0.045; Supplementary Table 15).

**Figure 3.**
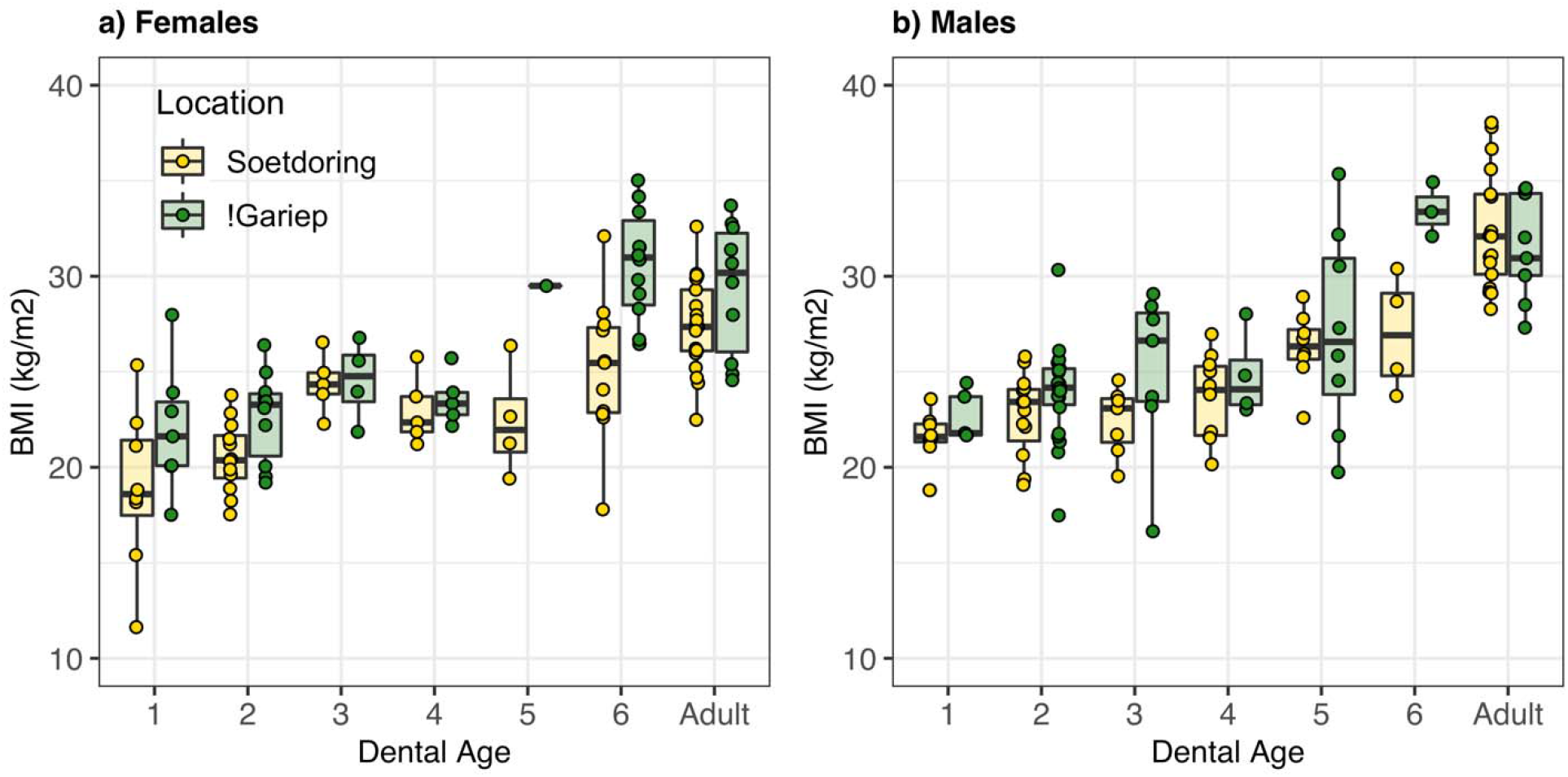
Body mass index or BMI (kg/m^2^) in a) female and b) male vervet monkeys across each dental age category. For estimated chronological ages, see Table 1.

**Figure 4.**
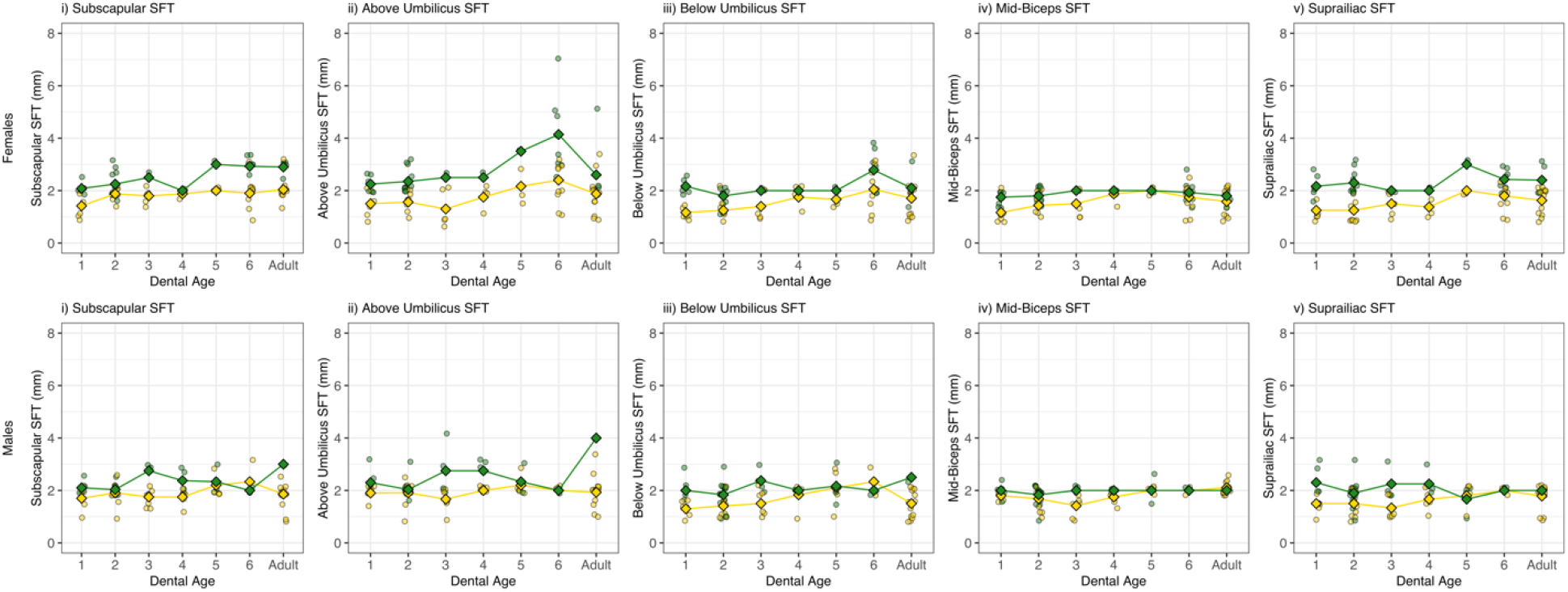
Mean i) subscapular, ii) above umbilicus, iii) below umbilicus, iv) mid-biceps, and v) suprailiac skin fold thickness in a) females, and b) males for vervets sampled in !Gariep (green) and Soetdoring (gold). Individual measurements are shown to demonstrate range.

### Male maturation – Relative testis volume

Differing patterns of testis growth distinguished males in Soetdoring from those in !Gariep, using both absolute and relative testis volume as measures (Fig 5). Although both absolute and relative testis volume appears larger in !Gariep for age categories 5 and 6, these differences were not significant. In adulthood, both relative and absolute testis volume was the same between the two sites.

**Figure 5.**
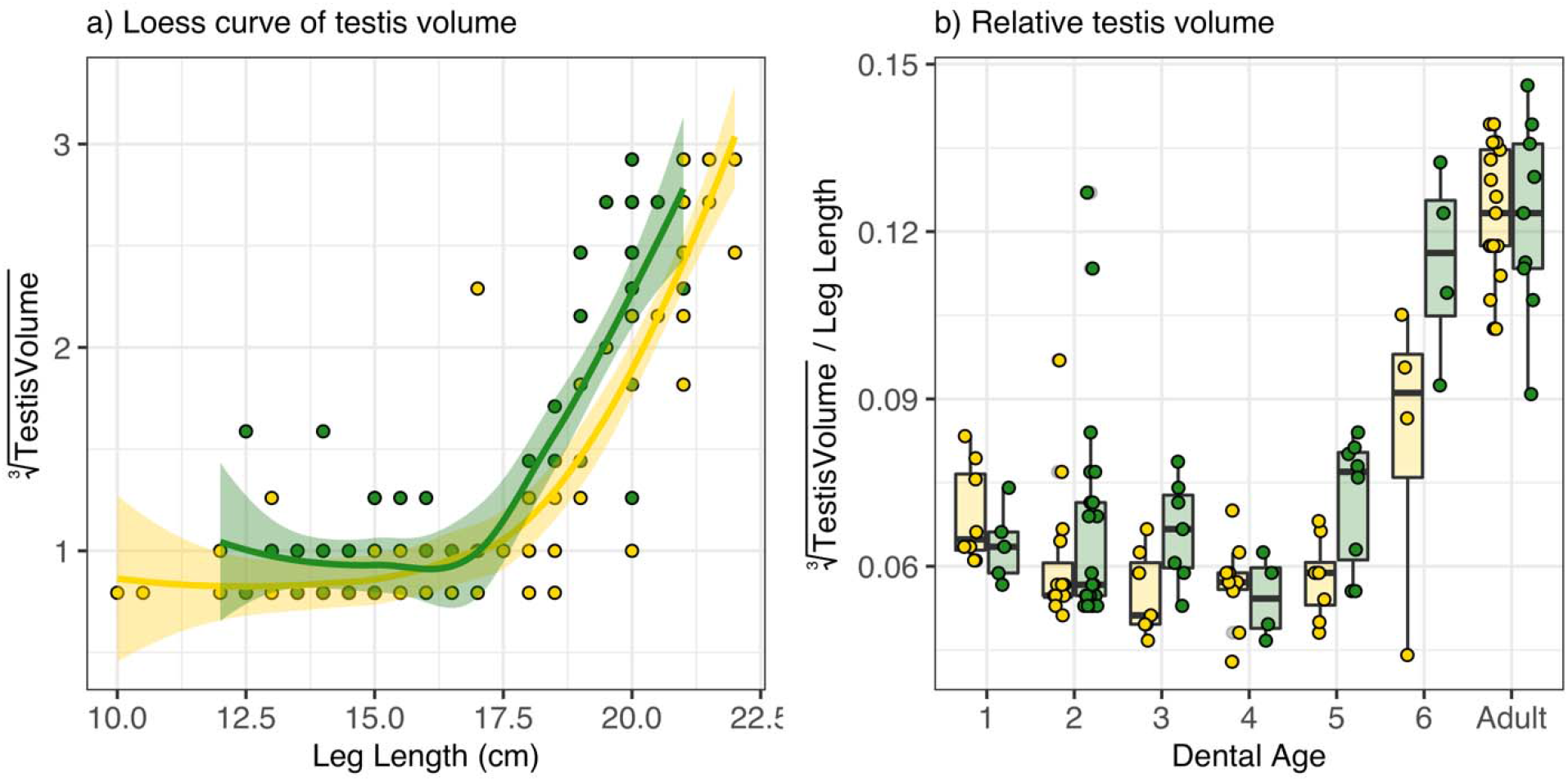
Boxplot of a) loess curves representing growth in absolute testis volume by body mass, and b) relative testis volume in males (testis volume/leg length) across each dental age category. See Table 2 for sample sizes. Gold indicates males sampled at Soetdoring Nature Reserve, while green represents males sampled in the !Gariep Dam region.

The pre-pubertal model with the lowest AICc included only body mass as a significant negative covariate of relative testis volume, albeit of small effect (β = −0.042, P = 0.0048; *R^2^* = 0.12; Table 3a). The peri-pubertal model with the lowest AICc included location (Soetdoring vs. !Gariep), dental age category, body mass, and an interaction term between age and body mass as covariates (Table 3b). In this model, males in !Gariep had significantly larger relative testis volume overall compared to those in Soetdoring (β = 0.088, P = 0.019; *R^2^* = 88.47). After accounting for body mass, dental age category was not significantly related to relative testis volume. Males in age category 6, however, showed a significantly larger increase in relative testis size as body mass increased (β = 0.295, P = 0.008).

**Table 3:** Models of Relative Testis Volume

| **a) Pre-Pubertal (Age Categories 1 - 4) Model** |  |  |  |  |
| --- | --- | --- | --- | --- |
| RTV ~ Body Mass |  |  |  |  |
|  | **Estimate** | **Error** | **p value** |  |
| **Intercept** | **-2.673** | **0.059** | **< 0.001** | **\*\*\*** |
| **Body Mass** | **-0.081** | **0.026** | **0.003** | **\*\*** |
| **b) Peri-Pubertal (Age Categories 4 - Adult) Model** |  |  |  |  |
| RTV ~ Age * Body Mass + Location |  |  |  |  |
|  | **Estimate** | **Error** | **p value** |  |
| **Intercept** | **-2.820** | **0.279** | **< 0.001** | **\*\*\*** |
| **Location (!Gariep)** | **0.088** | **0.037** | **0.019** | **\*** |
| **Age Category** | - | - | - |  |
| 5 | 0.038 | 0.334 | 0.909 |  |
| **6** | **-0.842** | **0.377** | **0.029** | **\*** |
| Adult | 0.021 | 0.363 | 0.954 |  |
| Body Mass | -0.029 | 0.094 | 0.760 |  |
| Body Mass : Age Category 5 | 0.028 | 0.107 | 0.797 |  |
| **Body Mass : Age Category 6** | **0.295** | **0.107** | **0.008** | **\*\*** |
| Body Mass : Age Category Adult | 0.140 | 0.102 | 0.174 |  |

### Female maturation – Parity status by nipple length

As indicated by nipple morphology, no female sampled in Soetdoring showed signs of parity before dental age 6 (Fig 6). Two Soetdoring females in the adult sample were primiparous. In !Gariep, primiparity began at dental age 4 (26-31 months), and eight females were already multiparous by dental age 6. All adult females (N = 30) had given birth at least once. The model with the lowest AICc for parity included body mass and location as covariates, along with their interaction term (*R^2^*=0.796; Table 4). Body mass, overall, had a significant positive association with parity status (β = 0.745, P = 0.009). No Fisher’s exact test of parity status across sites within age categories showed significant differences.

**Figure 6.**
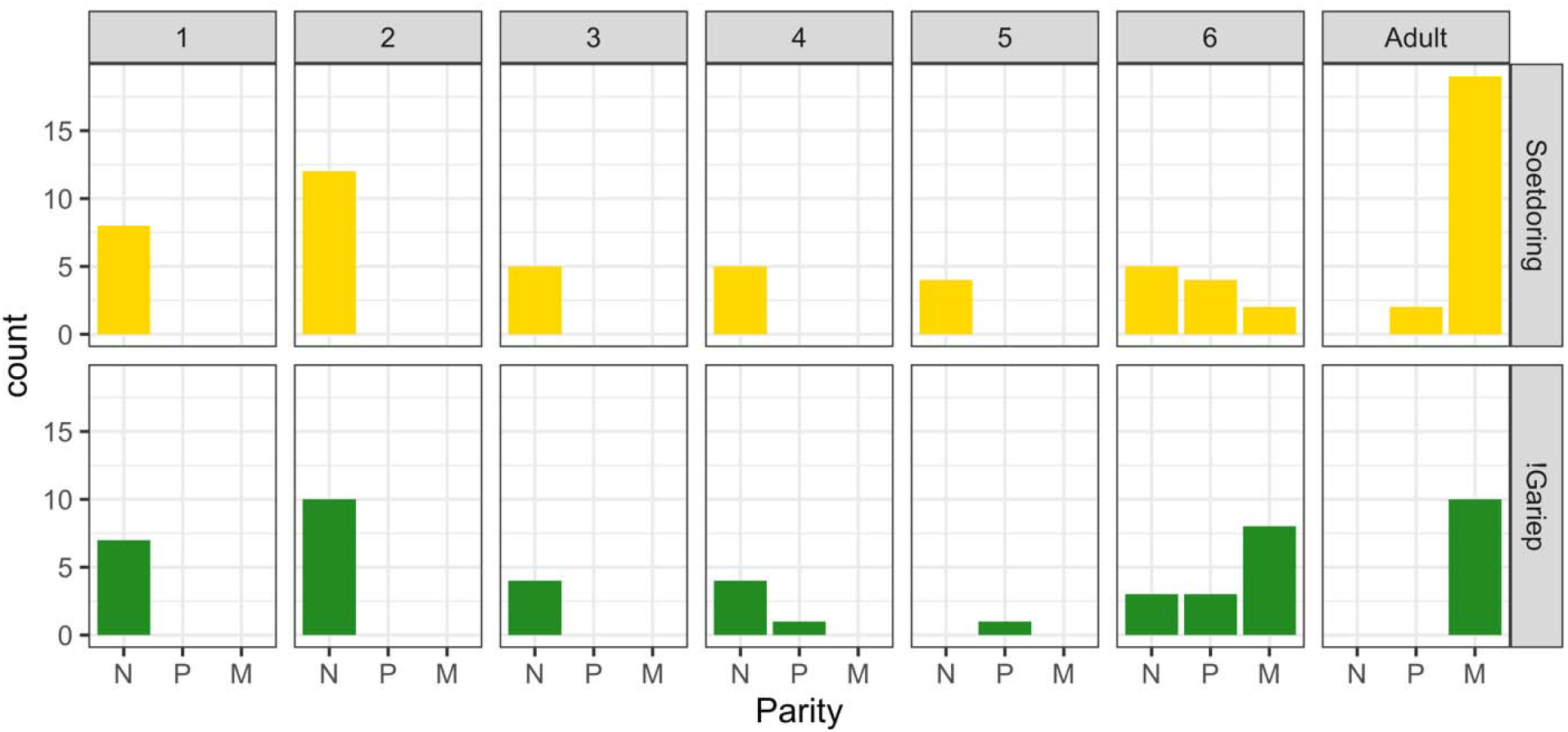
The number of females exhibiting nipple morphology indicative of each parity category (N = Nulliparous, P = Primiparous, M = Multiparous) within each dental age, compared between sites.

**Table 4:** Logistic Regression Model for Female Parity

| Parity ~ Body Mass*Location |  |  |  |  |
| --- | --- | --- | --- | --- |
|  | Estimate | Error | p value |  |
| Intercept | -23.482 | 9.094 | 0.010 | ** |
| Body Mass | 7.445 | 2.857 | 0.009 | ** |
| Location (!Gariep) | 11.486 | 9.736 | 0.238 |  |
| Body Mass : |  |  |  |  |
| Location(!Gariep) | -3.900 | 3.020 | 0.197 |  |

## DISCUSSION

The vervets in !Gariep show clear evidence of better body condition, with significantly thicker skin folds in all parts of the body measured, with significantly higher body mass and BMI overall, as well as within specific age/sex categories. Differences in body mass between the two sites become apparent in males at dental age 6 (Fig 2), when !Gariep males appear heavier than in Soetdoring. This timing is generally consistent with the pubertal growth spurt for males of this taxon (Turner et al. 2019; Jarrett et al. 2020). Given that males in !Gariep are not then significantly heavier in adulthood than those in Soetdoring, we interpret as evidence of an earlier peak of growth at !Gariep. Turner et al. (2019) note a typical pubertal growth spurt in female vervets in dental age 4 or 5 (after approximately 24 months), consistent with the generally earlier onset of reproduction in cercopithecine females relative to males (Bercovitch 2000). Although our sample size for these age categories is unfortunately small, we do note that the one !Gariep female sampled at dental age 5 was comparable in weight to dental age 6 females, and females remain significantly heavier in !Gariep compared to Soetdoring from this age through adulthood.

Given the earlier and more exaggerated increases in mass at !Gariep, it is unclear why adult males do not also appear heavier in !Gariep. That much larger male body size in provisioned groups appears to be the norm in previously studied populations (e.g., Altmann & Alberts 2005) makes this more puzzling. This could potentially be explained by male-biased dispersal in vervets, particularly in the context of these study groups (Henzi and Lucas 1980; Cheney and Seyfarth 1983; Turner et al. 2019). Provisioning is rare for the vervets in Soetdoring Nature Reserve, but the reserve is bordered entirely by cropland (Fig 1b). No one has observed the entire study groups sampled here crossing the Modder River into these agricultural areas, although on two separate occasions individuals, including an identified subadult male, were observed to swim across the river and back (Blaszczyk 2016; Nick Theron, personal communication). As such, although males may cross the Modder River on occasion and for dispersal purposes, the river may serve as a barrier limiting regular access to anthropogenic foods for females. Adult males that disperse into our study groups from these farms may have benefitted from consistent food enhancement from crops during development. Conversely, the farms in !Gariep are relatively isolated agricultural areas surrounded by bushveld and nature reserves (Fig 1c), suggesting that adult males dispersing into these farms may come from less well-provisioned areas. Given this, it is possible that we are seeing heavier than expected males in Soetdoring, and lighter than expected males in !Gariep, given the nutritional environments to which pre- and peri-pubertal males in these populations have access.

Our best model for pre-pubertal testis size suggests a significant decrease in relative testis size with age until puberty. This result reflects a relative stasis in testis size as body mass increases through the juvenile phase of development. Along with an earlier significant increase in body mass, peri-pubertal males in !Gariep show significantly larger relative testis volume than in Soetdoring in dental ages 5 and 6 (Fig 3), ultimately leading to similar relative testis volumes by adulthood. Increases in testosterone levels accompanied by a 15% increase in testis volume in captive vervets during the breeding season suggest that a larger testis volume may confer a higher chance of fertilization (Eley et al. 1986; Eley 1992). Sample collection was limited to June through August, which is just after the peak breeding season, characterized as April to June throughout South Africa (Blaszczyk 2016; McFarland et al. 2014). Given that our sampling was constrained to this season alone, we do not think that seasonal effects are a concern. Intraspecific variation in testis size is associated with higher circulating testosterone concentration, sperm quality, and fitness (Schulte-Hostedde et al. 2004; Hamada et al. 2005). However, the larger testis volume during these ages in !Gariep appears to be a developmental pattern, not leading to substantial adult differences translatable to long-term increases in fertility beyond a potentially earlier initiation of reproduction. This earlier reproductive maturation, if paired with an earlier age at dispersal, could lead to an earlier age at first reproduction in males and higher fitness. Previous work in Soetdoring, however, only observed the intergroup transfer of adults and one very large subadult male (Blasczyzk 2016); we have not yet carried out long term behavioral observations in !Gariep populations. To assess whether the earlier increase in testis volume in !Gariep could lead to earlier reproduction will require more extensive behavioral observations and direct continuous monitoring of these populations.

Despite evidence of primiparity at dental ages 4 and 5 in !Gariep, our sample size limits our ability to interpret the relative timing of reproductive onset in females between these populations. Given that nipple morphology appears to be directly related to use, this suggests that female vervets begin nursing at a younger age on the !Gariep farms, indicating earlier onset of reproduction. Additionally, nearly half of the females in dental age 6 in !Gariep were observed to be lactating, compared to none in Soetdoring (although nearly half of the females in this age category were noted to be pregnant in both populations). Given these indicators are reliable signs of earlier reproduction in !Gariep, it would be consistent with the human and non-human primate literature showing strong correlations between nutritional enrichment and earlier menarche and age at first birth (Mori 1979; Cheney et al. 1988; Altmann and Alberts 2005; Gluckman and Hanson 2006a), having reached a critical threshold of body fatness or body mass for reproductive viability at an earlier age (e.g., Wade & Schneider 1996). The overall higher body mass and BMI of pregnant and lactating females in both dental age 6 and adulthood could reflect that pattern, and suggest that the earlier attainment of heavier body mass allowed !Gariep females to initiate reproduction sooner. Alternatively, the added weight and BMI could be a byproduct of their reproductive status, reflecting weight-gain associated with pregnancy or in preparation for extended lactation (McFarland 1997).

One weakness of this study is that we lack longitudinal behavioral observations of foraging/feeding, copulation, and birth to pair with these proxy measures. Calorie-rich crop foraging is a logical conclusion given the otherwise uniform natural ecologies of these sites. Still, without foraging observations we cannot demonstrate which environmental factor at !Gariep is directly responsible for the earlier attainment of these maturational landmarks. We were also unable to control for behavioral factors that may have influenced access to anthropogenic resources, including rank. In wild populations, rank mediates priority of access to resources, leading to accelerated growth and earlier maturation in more highly ranked individuals (Whitten 1983; Bercovitch & Strum 1993; Onyango et al., 2013; Jarrett et al., 2020). This same priority of access influences access to anthropogenic food resources in urban macaques, with males and high ranking females getting more caloric benefits, potentially limiting the fitness benefits of these foods for lower-ranking females (Marty et al. 2019).

Additionally, without hormonal data it is difficult to directly link the evidently higher body fatness in !Gariep with the apparently earlier onset of reproduction. In many cases of early reproductive onset in humans and non-human primates, increased caloric intake is thought to drive this pattern (Altmann and Alberts 2005; Gluckman and Hanson 2006a; Wade & Schneider 1996). However, there are known mediating factors. Low birth weight derived from insufficient nutrition *in utero*, and rapid postnatal growth during critical developmental stages, for example, often precede early reproductive maturation in human and non-human primates of both sexes (Ibáñez et al. 2000; Kuzawa et al. 2010). High age-specific levels of body fatness and associated hormone levels, such as elevated circulating levels of the adipose-derived hormone leptin, are also associated with accelerated developmental trajectories and the earlier timing of reproductive maturation (Bercovitch 2000; Whitten & Turner 2009; Gluckman and Hanson 2006b). An assessment of circulating leptin levels and sex steroids would make the link between body fatness and early reproductive onset more clear. Future research should also consider other environmental factors that could alter developmental constraints on growth and secondary sexual characteristics, including potential anthropogenic determinants like endocrine-disrupting agrochemicals (e.g., English et al. 2012; Blanck et al. 2000).

While we have focused on the benefits of anthropogenic food-enhancement in developmental timing in these populations, the costs of human-wildlife conflict may also be affecting life history and development in !Gariep. Vervets are classified as ‘vermin’ in South Africa, and permits are not required to kill them on private property. We have not observed attempts to capture or harm vervet monkeys on the farms in !Gariep first-hand, but several monkeys present with missing legs (indicating past encounters with snares), evidence of electrocution (presumably by power lines), and metal BBs under their skin (suggesting their having been shot by humans). Although the owners of each farm in !Gariep Dam, by all accounts, treat the resident monkeys quite well, this does not preclude conflict on other lands. Indeed, most evidence of these risks is seen in adult males, who no doubt emigrated into our study groups. High psychosocial stress is associated with delays in growth and reproductive onset (Johnson 2003; Onyango et al., 2013), but such unpredictable and dangerous environments are also associated with an earlier onset of menarche and first birth (Chisolm et al. 2005; Ellis et al. 2009). A closer examination of individual risks and stressors in each landscape would be required to tease out what role, if any, the stressors of each environment play independently of nutrition in the timing of reproductive onset.

Addressing potential demographic effects on the timing of maturation was beyond the scope of this project, but should be considered in studies with longitudinal or more diverse population samples. In Amboseli baboons, relatively small populations appear to experience earlier maturation, presumably due to lower competition for resources (Altmann & Alberts 2003). While the !Gariep populations are larger and occur at a higher density than those in Soetdoring (Schmitt, unpublished data), this does not appear to limit growth and maturation. This may be because demography has not yet outstripped the available food resources for these monkeys. However, high population density and the hypothesized risks of high mortality environments are thought to occur hand-in-hand (Ellis et al. 2009). Such populations may then also experience selection for faster life histories with earlier reproductive maturity (Ellis et al. 2009; Wells 2012). A better understanding of the risks and stresses faced by each population is necessary to clarify how these factors may also contribute to the timing of maturation.

This work demonstrates a potential effect of anthropogenic food enhancement on the reproductive maturation among wild vervet monkeys, underscoring previous research on this topic. Our results add to the evidence indicating an increase in body condition and more rapid reproduction for primates living in anthropogenic environments. Future research on these populations should add to these results with detailed behavioral and nutritional data, more detailed data on ecological stressors, and physiological indicators of energy balance and maturation. The continued incursion of human environments on non-human primate habitats around the world (Estrada et al. 2017) demand increased attention to the effects of human presence and resources on primate biology and health.

## ACKNOWLEDGEMENTS

We thank our collaborator Adrian Tordiffe, at the University of Pretoria Faculty of Veterinary Medicine, for his logistical help and aid with permits. The staff of the Soetdoring Nature Reserve have been very helpful, most especially Contance Simwinji. Our work at Southford Stud was made possible by the generosity and hospitality of David and Kathleen Southey, and at Shekinah/Orange Valley by Eben Olivier, Candyce and Louis Fritz, and Careena Taylor. We would like to thank the Free State Department of Economic Development, Tourism, and Environmental Affairs, without whom this research would not have been possible, particularly the efforts of Nacelle Collins and Johannes Mosia. We thank the hard work of all the veterinarians who have assisted with data collection and safeguarded the well-being of the vervets while in our care, including Lizanne Meiring, Gavin and Charmaine Rous, and most especially Rudolf Venter of the Motsumi Dierikliniek. We thank Morgan Farrar, Christian Gagnon, Sara Jablonski, Sasami Langford, Evan Razdan, and Amy Scott for help with data collection in 2016-2018, as well as Liza and Krona at Truksvy for logistical support. Initial collections of this data in 2010 would not have been possible without the efforts of Anna Jasinska, Yoon Jung, Gabriel Coetzer, Elzet Aswegen, Tegan Gaetano, Dewald DuPlessis, Micah Beller, David Beller, Murray Stokoe, and Oliver “Pess” Morton. Helpful conversations and suggestions regarding this study from the 2019 meeting of the Primate Ecology and Genetics Group (the South African Primatological Society) at the National Zoological Gardens in Pretoria helped this work along immensely. This research was accomplished with all applicable permissions in both the US and South Africa, including from the Free State Department of Economic Development, Tourism and Environmental Affairs; the Northern Cape Department of Environment and Nature Conservation; the South Africa Department of Agriculture, Forestry & Fisheries; and the South African National Department of Environmental Affairs. All procedures involving the capture, sample collection, and handling of live non-human primates were approved by the Boston University Institutional Animal Care and Use Committee; University of Wisconsin, Milwaukee IACUC; the UCLA Animal Research Committee; and University of the Free State Animal Research Ethics Committee.

