## Supplementary Table for "Anthropogenic food enhancement alters the timing of maturational landmarks among wild savanna monkeys (*Chlorocebus pygerythrus*)"

| **Supplementary Table 2: Body Mass and Pregnancy Status** | | | | |  | |  | |  | |  | |  | |
| --- | --- | --- | --- | --- | --- | --- | --- | --- | --- | --- | --- | --- | --- | --- |
| **a) Type-II Welch ANOVA for Pregnancy Status and Body Mass** | | | | | | |  | |  | |  | |  | |
| Body Mass ~ Pregnant*Age*Location | |  | |  |  | |  | |  | |  | |  | |
|  |  | **F** (5,31) | | **df** | **p value** | |  | |  | |  | |  | |
| **Pregnant** | | **5.40** | | **1** | **0.027** | | * | |  | |  | |  | |
| **Age Category** | | **19.67** | | **1** | **< 0.001** | | *** | |  | |  | |  | |
| **Location** | | **32.27** | | **1** | **< 0.001** | | *** | |  | |  | |  | |
| Pregnant:Age Category | | 0.01 | | 1 | 0.941 | |  | |  | |  | |  | |
| Location:Age Category | | 0.00 | | 1 | 0.982 | |  | |  | |  | |  | |
| Pregnant:Location | | 0.07 | | 1 | 0.791 | |  | |  | |  | |  | |
| Pregnant:Location:Age Category | | 0.67 | | 1 | 0.418 | |  | |  | |  | |  | |
| **b) Games-Howell Post Hoc Comparisons** | | | |  |  | |  | |  | |  | |  | |
|  | | **Pregnant** | | **n** | **Marginal Means** | | **Estimate** | | **df** | | **t** | | **p value** | |
| Soetdoring Age 6 | | 0 | | 4 | 3.28 (2.88,3.68) | | 0.25 | | 7.10 | | 1.06 | | 0.948 | |
|  |  | 1 | | 6 | 3.53 (3.20,3.86) | |  |  |  |  |  |  |  |  |
| Soetdoring Adult | | 0 | | 10 | 3.75 (3.50,4.01) | | 0.53 | | 11.94 | | -1.03 | | NA | |
|  |  | 1 | | 2 | 4.29 (3.72,4.85) | |  |  |  |  |  |  |  |  |
| !Gariep Age 6 | | 0 | | 6 | 3.89 (3.56,4.21) | | 0.47 | | 7.30 | | 2.15 | | 0.465 | |
|  |  | 1 | | 5 | 4.35 (4.00,4.71) | |  |  |  |  |  |  |  |  |
| !Gariep Adult | | 0 | | 4 | 4.45 (4.05,4.85) | | 0.04 | | 1.04 | | 0.09 | | NA | |
|  |  | 1 | | 2 | 4.50 (3.93,5.06) | |  |  |  |  |  |  |  |  |
| **Supplementary Table 3: BMI and Pregnancy Status** | | | | |  | |  | |  | |  | |  | |
| **a) Type-II Welch ANOVA for Pregnancy Status and BMI** | | | | |  | |  | |  | |  | |  | |
| BMI ~ Pregnant*Age*Location | |  | |  |  | |  | |  | |  | |  | |
|  |  | **F** (267,31) | | **df** | **p value** | |  | |  | |  | |  | |
| **Pregnant** | | **4.64** | | **1** | **0.039** | | * | |  | |  | |  | |
| Age Category | | 2.31 | | 1 | 0.139 | |  | |  | |  | |  | |
| **Location** | | **17.63** | | **1** | **< 0.001** | | *** | |  | |  | |  | |
| Pregnant:Age Category | | 0.01 | | 1 | 0.929 | |  | |  | |  | |  | |
| Location:Age Category | | 1.76 | | 1 | 0.195 | |  | |  | |  | |  | |
| Pregnant:Location | | 0.80 | | 1 | 0.378 | |  | |  | |  | |  | |
| Pregnant:Location:Age Category | | 0.04 | | 1 | 0.852 | |  | |  | |  | |  | |
| **b) Games-Howell Post Hoc Comparisons** | | | |  |  | |  | |  | |  | |  | |
|  | | **Pregnant** | | **n** | **Marginal Means** | | **Estimate** | | **df** | | **t** | | **p value** | |
| Soetdoring Age 6 | | 0 | | 4 | 24.2 (21.2,27.2) | | 2.65 | | 7.85 | | 1.56 | | 0.758 | |
|  |  | 1 | | 6 | 26.9 (24.4,29.3) | |  |  |  |  |  |  |  |  |
| Soetdoring Adult | | 0 | | 10 | 26.6 (24.7,28.5) | | 2.68 | | 4.08 | | 2.27 | | 0.456 | |
|  |  | 1 | | 2 | 29.2 (25.0,33.5) | |  |  |  |  |  |  |  |  |
| !Gariep Age 6 | | 0 | | 6 | 30.8 (28.3,33.2) | | 0.90 | | 8.75 | | 0.50 | | 0.999 | |
|  |  | 1 | | 5 | 31.7 (29.0,34.3) | |  |  |  |  |  |  |  |  |
| !Gariep Adult | | 0 | | 4 | 30.3 (27.3,33.3) | | 0.01 | | 2.36 | | 0.00 | | 1.000 | |
|  |  | 1 | | 2 | 30.3 (26.0,34.5) | |  |  |  |  |  |  |  |  |
| **Supplementary Table 4: Above Umbilicus SFT and Pregnancy Status** | | | | | | |  | |  | |  | |  | |
| **a) Type-II Welch ANOVA for Pregnancy Status and SFT Above Umbilicus** | | | | | | |  | |  | |  | |  | |
| SFT AU ~ Pregnant*Age*Location | |  | |  |  | |  | |  | |  | |  | |
|  |  | **F** (34,27) | | **df** | **p value** | |  | |  | |  | |  | |
| Pregnant | | 0.58 | | 1 | 0.452 | |  | |  | |  | |  | |
| Age Category | | 2.04 | | 1 | 0.164 | |  | |  | |  | |  | |
| Location | | 2.80 | | 1 | 0.106 | |  | |  | |  | |  | |
| Pregnant:Age Category | | 0.48 | | 1 | 0.494 | |  | |  | |  | |  | |
| Location:Age Category | | 1.63 | | 1 | 0.213 | |  | |  | |  | |  | |
| Pregnant:Location | | 1.52 | | 1 | 0.228 | |  | |  | |  | |  | |
| Pregnant:Location:Age Category | | 1.75 | | 1 | 0.197 | |  | |  | |  | |  | |
| **b) Games-Howell Post Hoc Comparisons** | | | |  |  | |  | |  | |  | |  | |
|  | | **Pregnant** | | **n** | **Marginal Means** | | **Estimate** | | **df** | | **t** | | **p value** | |
| Soetdoring Age 6 | | 0 | | 4 | 2.00 (0.85,3.15) | | 0.67 | | 7.61 | | 1.14 | | 0.929 | |
|  |  | 1 | | 6 | 2.67 (1.73,3.60) | |  |  |  |  |  |  |  |  |
| Soetdoring Adult | | 0 | | 10 | 1.75 (1.02,2.48) | | 0.75 | | 1.50 | | 1.35 | | NA | |
|  |  | 1 | | 2 | 2.50 (0.88,4.12) | |  |  |  |  |  |  |  |  |
| !Gariep Age 6 | | 0 | | 3 | 3.50 (2.17,4.83) | | 1.13 | | 4.99 | | 0.95 | | 0.964 | |
|  |  | 1 | | 4 | 4.62 (3.48,5.77) | |  |  |  |  |  |  |  |  |
| !Gariep Adult | | 0 | | 4 | 3.38 (2.23,4.52) | | -1.38 | | 3.00 | | -1.72 | | 1.000 | |
|  |  | 1 | | 2 | 2.00 (0.38,3.62) | |  |  |  |  |  |  |  |  |
| **Supplementary Table 5: Below Umbilicus SFT and Pregnancy Status** | | | | | | |  | |  | |  | |  | |
| **a) Type-II Welch ANOVA for Pregnancy Status and SFT Below Umbilicus** | | | | | | |  | |  | |  | |  | |
| SFT BU ~ Pregnant*Age*Location | |  | |  |  | |  | |  | |  | |  | |
|  |  | **F** (15,27) | | **df** | **p value** | |  | |  | |  | |  | |
| Pregnant | | 0.30 | | 1 | 0.586 | |  | |  | |  | |  | |
| Age Category | | 2.31 | | 1 | 0.141 | |  | |  | |  | |  | |
| Location | | 1.27 | | 1 | 0.269 | |  | |  | |  | |  | |
| Pregnant:Age Category | | 0.00 | | 1 | 0.999 | |  | |  | |  | |  | |
| Location:Age Category | | 1.52 | | 1 | 0.229 | |  | |  | |  | |  | |
| Pregnant:Location | | 2.96 | | 1 | 0.097 | |  | |  | |  | |  | |
| Pregnant:Location:Age Category | | 0.03 | | 1 | 0.874 | |  | |  | |  | |  | |
| **b) Games-Howell Post Hoc Comparisons** | | | |  |  | |  | |  | |  | |  | |
|  | | **Pregnant** | | **n** | **Marginal Means** | | **Estimate** | | **df** | | **t** | | **p value** | |
| Soetdoring Age 6 | | 0 | | 4 | 1.88 (1.11,2.64) | | 0.29 | | 5.50 | | 0.57 | | 0.998 | |
|  |  | 1 | | 6 | 2.17 (1.54,2.79) | |  |  |  |  |  |  |  |  |
| Soetdoring Adult | | 0 | | 10 | 1.65 (1.17,2.14) | | 0.35 | | 9.00 | | 1.35 | | 0.857 | |
|  |  | 1 | | 2 | 2.00 (0.92,3.09) | |  |  |  |  |  |  |  |  |
| !Gariep Age 6 | | 0 | | 3 | 3.00 (2.11,3.89) | | -0.38 | | 3.61 | | -0.54 | | 1.000 | |
|  |  | 1 | | 4 | 2.62 (1.86,3.39) | |  |  |  |  |  |  |  |  |
| !Gariep Adult | | 0 | | 4 | 2.25 (1.48,3.02) | | -0.50 | | 3.00 | | -1.41 | | 1.000 | |
|  |  | 1 | | 2 | 1.75 (0.67,2.84) | |  |  |  |  |  |  |  |  |
| **Supplementary Table 6: Subscapular SFT and Pregnancy Status** | | | | | | |  | |  | |  | |  | |
| **a) Type-II Welch ANOVA for Pregnancy Status and SFT Subscapular** | | | | | | |  | |  | |  | |  | |
| SFT Subscap ~ Pregnant*Age*Location | |  | |  |  | |  | |  | |  | |  | |
|  |  | **F** (4,27) | | **df** | **p value** | |  | |  | |  | |  | |
| Pregnant | | 0.11 | | 1 | 0.737 | |  | |  | |  | |  | |
| Age Category | | 0.00 | | 1 | 0.973 | |  | |  | |  | |  | |
| **Location** | | **54.67** | | **1** | **< 0.001** | | *** | |  | |  | |  | |
| Pregnant:Age Category | | 0.23 | | 1 | 0.634 | |  | |  | |  | |  | |
| Location:Age Category | | 2.86 | | 1 | 0.103 | |  | |  | |  | |  | |
| Pregnant:Location | | 3.17 | | 1 | 0.086 | |  | |  | |  | |  | |
| Pregnant:Location:Age Category | | 0.04 | | 1 | 0.839 | |  | |  | |  | |  | |
| **b) Games-Howell Post Hoc Comparisons** | | | |  |  | |  | |  | |  | |  | |
|  | | **Pregnant** | | **n** | **Marginal Means** | | **Estimate** | | **df** | | **t** | | **p value** | |
| Soetdoring Age 6 | | 0 | | 4 | 1.62 (1.21,2.04) | | 0.46 | | 6.71 | | 1.47 | | 0.803 | |
|  |  | 1 | | 6 | 2.08 (1.75,2.42) | |  |  |  |  |  |  |  |  |
| Soetdoring Adult | | 0 | | 10 | 1.95 (1.69,2.21) | | 0.55 | | 1.02 | | 1.09 | | NA | |
|  |  | 1 | | 2 | 2.50 (1.92,3.08) | |  |  |  |  |  |  |  |  |
| !Gariep Age 6 | | 0 | | 3 | 3.17 (2.69,3.64) | | -0.42 | | 4.35 | | -1.15 | | 1.000 | |
|  |  | 1 | | 4 | 2.75 (2.34,3.16) | |  |  |  |  |  |  |  |  |
| !Gariep Adult | | 0 | | 4 | 3.88 (2.46,3.29) | | -0.13 | | 1.53 | | -0.45 | | 1.000 | |
|  |  | 1 | | 2 | 2.75 (2.17,3.33) | |  |  |  |  |  |  |  |  |
| **Supplementary Table 7: Mid-Biceps SFT and Pregnancy Status** | | | | | | |  | |  | |  | |  | |
| **a) Type-II Welch ANOVA for Pregnancy Status and SFT Mid-Biceps** | | | | | | |  | |  | |  | |  | |
| SFT MB ~ Pregnant*Age*Location | |  | |  |  | |  | |  | |  | |  | |
|  |  | **F** (6,27) | | **df** | **p value** | |  | |  | |  | |  | |
| **Pregnant** | | **10.00** | | **1** | **0.004** | | ******* | |  | |  | |  | |
| Age Category | | 1.36 | | 1 | 0.255 | |  | |  | |  | |  | |
| Location | | 0.00 | | 1 | 1.000 | |  | |  | |  | |  | |
| Pregnant:Age Category | | 1.55 | | 1 | 0.224 | |  | |  | |  | |  | |
| Location:Age Category | | 0.01 | | 1 | 0.931 | |  | |  | |  | |  | |
| Pregnant:Location | | 1.78 | | 1 | 0.193 | |  | |  | |  | |  | |
| Pregnant:Location:Age Category | | 0.28 | | 1 | 0.601 | |  | |  | |  | |  | |
| **b) Games-Howell Post Hoc Comparisons** | | | |  |  | |  | |  | |  | |  | |
|  | | **Pregnant** | | **n** | **Marginal Means** | | **Estimate** | | **df** | | **t** | | **p value** | |
| Soetdoring Age 6 | | 0 | | 4 | 1.62 (1.15,2.10) | | 0.21 | | 7.00 | | 0.65 | | 1.000 | |
|  |  | 1 | | 6 | 1.83 (1.45,2.22) | |  |  |  |  |  |  |  |  |
| Soetdoring Adult | | 0 | | 10 | 1.50 (1.20,1.80) | | 0.50 | | 9.00 | | 2.25 | | 1.000 | |
|  |  | 1 | | 2 | 2.00 (1.33,2.67) | |  |  |  |  |  |  |  |  |
| !Gariep Age 6 | | 0 | | 3 | 2.17 (1.62,2.71) | | -0.42 | | 2.40 | | 0.90 | | 0.960 | |
|  |  | 1 | | 4 | 1.75 (1.28,2.22) | |  |  |  |  |  |  |  |  |
| !Gariep Adult | | 0 | | 4 | 1.75 (1.28,2.22) | | 0.25 | | 3.00 | | 1.73 | | 0.690 | |
|  |  | 1 | | 2 | 2.00 (1.33,2.67) | |  |  |  |  |  |  |  |  |
| **Supplementary Table 8: Suprailiac SFT and Pregnancy Status** | | | | | | |  | |  | |  | |  | |
| **a) Type-II Welch ANOVA for Pregnancy Status and SFT Suprailiac** | | | | | | |  | |  | |  | |  | |
| SFT Suprailiac ~ Pregnant*Age*Location | |  | |  |  | |  | |  | |  | |  | |
|  |  | **F** (6,27) | | **df** | **p value** | |  | |  | |  | |  | |
| **Pregnant** | | **4.12** | | **1** | **0.052** | | * | |  | |  | |  | |
| Age Category | | 0.40 | | 1 | 0.531 | |  | |  | |  | |  | |
| Location | | 0.00 | | 1 | 1.000 | |  | |  | |  | |  | |
| Pregnant:Age Category | | 0.24 | | 1 | 0.631 | |  | |  | |  | |  | |
| Location:Age Category | | 0.81 | | 1 | 0.377 | |  | |  | |  | |  | |
| **Pregnant:Location** | | **7.51** | | **1** | **0.011** | | ** | |  | |  | |  | |
| Pregnant:Location:Age Category | | 0.23 | | 1 | 0.634 | |  | |  | |  | |  | |
| **b) Games-Howell Post Hoc Comparisons** | | | |  |  | |  | |  | |  | |  | |
|  | | **Pregnant** | | **n** | **Marginal Means** | | **Estimate** | | **df** | | **t** | | **p value** | |
| Soetdoring Age 6 | | 0 | | 4 | 1.50 (1.00,2.00) | | 0.21 | | 6.95 | | 0.65 | | 1.000 | |
|  |  | 1 | | 6 | 2.00 (1.59,2.41) | |  |  |  |  |  |  |  |  |
| Soetdoring Adult | | 0 | | 10 | 1.55 (1.23,1.87) | | 0.50 | | 9.00 | | 3.35 | | 0.100 | |
|  |  | 1 | | 2 | 2.00 (1.29,2.71) | |  |  |  |  |  |  |  |  |
| !Gariep Age 6 | | 0 | | 3 | 2.50 (1.92,3.08) | | -0.42 | | 2.43 | | 0.90 | | 0.960 | |
|  |  | 1 | | 4 | 2.38 (1.88,2.87) | |  |  |  |  |  |  |  |  |
| !Gariep Adult | | 0 | | 4 | 2.50 (2.00,3.00) | | 0.25 | | 3.00 | | 1.73 | | 0.690 | |
|  |  | 1 | | 2 | 2.00 (1.29,2.71) | |  |  |  |  |  |  |  |  |
| **Supplementary Table 9: Body Mass and Lactation Status** | | |  | | |  | |  | |  | |  | |  |
| **a) Type-II ANOVA for Lactation Status and Body Mass** | | |  | | |  | |  | |  | |  | |  |
| Body Mass ~ Lactating*Age*Location |  | |  | | |  | |  | |  | |  | |  |
|  | **F** (9,47) | | **df** | | | **p value** | |  | |  | |  | |  |
| **Lactating** | **8.35** | | **1** | | | **0.006** | | ** | |  | |  | |  |
| **Age Category** | **13.22** | | **1** | | | **< 0.001** | | *** | |  | |  | |  |
| **Location** | **12.82** | | **1** | | | **< 0.001** | | *** | |  | |  | |  |
| Lactating:Age Category | 0.66 | | 1 | | | 0.422 | |  | |  | |  | |  |
| Location:Age Category | 0.31 | | 1 | | | 0.886 | |  | |  | |  | |  |
| Lactating:Location | 0.02 | | 1 | | | 0.580 | |  | |  | |  | |  |
| Lactating:Location:Age Category | - | | - | | | - | |  | |  | |  | |  |
| **b) Games-Howell Post Hoc Comparisons** |  | |  | | |  | |  | |  | |  | |  |
|  | **Lactating** | | **n** | | | **Marginal Means** | | **Estimate** | | **df** | | **t** | | **p value** |
| Soetdoring Age 6 | 0 | | 11 | | | 3.35 (3.09,3.61) | | - | | - | | - | | - |
|  | 1 | | 0 | | | - | |  |  |  |  |  |  |  |
| Soetdoring Adult | 0 | | 9 | | | 3.85 (3.56,4.13) | | 0.24 | | 17.48 | | 1.25 | | 0.867 |
|  | 1 | | 11 | | | 4.09 (3.83,4.35) | |  |  |  |  |  |  |  |
| !Gariep Age 6 | 0 | | 7 | | | 3.68 (3.35,4.00) | | 0.63 | | 11.98 | | 3.38 | | 0.061 |
|  | 1 | | 7 | | | 4.31 (3.99,4.64) | |  |  |  |  |  |  |  |
| !Gariep Adult | 0 | | 2 | | | 4.39 (3.79,5.00) | | 0.30 | | 5.99 | | 1.31 | | 0.828 |
|  | 1 | | 7 | | | 4.69 (4.37,5.02) | |  |  |  |  |  |  |  |
| **Supplementary Table 10: BMI and Lactation Status** | | |  | | |  | |  | |  | |  | |  |
| **a) Type-II ANOVA for Lactation Status and BMI** | | |  | | |  | |  | |  | |  | |  |
| BMI ~ Lactating*Age*Location |  | |  | | |  | |  | |  | |  | |  |
|  | **F** (263,47) | | **df** | | | **p value** | |  | |  | |  | |  |
| **Lactating** | **4.05** | | **1** | | | **0.050** | | * | |  | |  | |  |
| Age Category | 0.47 | | 1 | | | 0.495 | |  | |  | |  | |  |
| **Location** | **16.73** | | **1** | | | **< 0.001** | | *** | |  | |  | |  |
| Lactating:Age Category | 0.33 | | 1 | | | 0.571 | |  | |  | |  | |  |
| Location:Age Category | 2.17 | | 1 | | | 0.147 | |  | |  | |  | |  |
| Lactating:Location | 1.98 | | 1 | | | 0.166 | |  | |  | |  | |  |
| Lactating:Location:Age Category | - | | - | | | - | |  | |  | |  | |  |
| **b) Games-Howell Post Hoc Comparisons** |  | |  | | |  | |  | |  | |  | |  |
|  | **Lactating** | | **n** | | | **Marginal Means** | | **Estimate** | | **df** | | **t** | | **p value** |
| Soetdoring Age 6 | 0 | | 11 | | | 25.1 (23.4,26.8) | | - | | - | | - | | - |
|  | 1 | | 0 | | | - | |  |  |  |  |  |  |  |
| Soetdoring Adult | 0 | | 9 | | | 27.1 (25.2,28.9) | | 0.49 | | 17.77 | | 0.47 | | 0.999 |
|  | 1 | | 11 | | | 27.6 (25.9,29.2) | |  |  |  |  |  |  |  |
| !Gariep Age 6 | 0 | | **7** | | | 29.3 (27.2,31.4) | | 2.55 | | 10.13 | | 1.75 | | 0.605 |
|  | 1 | | **7** | | | 31.9 (29.8,34.0) | |  |  |  |  |  |  |  |
| !Gariep Adult | 0 | | 2 | | | 27.5 (23.6,31.5) | | 4.08 | | 1.23 | | 1.81 | | NA |
|  | 1 | | 7 | | | 31.6 (29.5,33.7) | |  |  |  |  |  |  |  |
| **Supplementary Table 11: Above Umbilicus SFT and Lactation Status** | | | | | |  | |  | |  | |  | |  |
| **a) Type-II ANOVA for Lactation Status and SFT Above Umbilicus** | | | | | |  | |  | |  | |  | |  |
| SFT AU ~ Lactating*Age*Location |  | |  | | |  | |  | |  | |  | |  |
|  | **F** (36,28) | | **df** | | | **p value** | |  | |  | |  | |  |
| Lactating | 1.29 | | 1 | | | 0.265 | |  | |  | |  | |  |
| Age Category | 3.81 | | 1 | | | 0.061 | |  | |  | |  | |  |
| **Location** | **5.41** | | **1** | | | **0.027** | | * | |  | |  | |  |
| Lactating:Age Category | 0.04 | | 1 | | | 0.849 | |  | |  | |  | |  |
| Location:Age Category | 0.84 | | 1 | | | 0.368 | |  | |  | |  | |  |
| Lactating:Location | 1.81 | | 1 | | | 0.189 | |  | |  | |  | |  |
| Lactating:Location:Age Category | - | | - | | | - | |  | |  | |  | |  |
| **b) Games-Howell Post Hoc Comparisons** |  | |  | | |  | |  | |  | |  | |  |
|  | **Lactating** | | **n** | | | **Marginal Means** | | **Estimate** | | **df** | | **t** | | **p value** |
| Soetdoring Age 6 | 0 | | 10 | | | 2.40 (1.70,3.13) | | - | | - | | - | | - |
|  | 1 | | 0 | | | - | |  |  |  |  |  |  |  |
| Soetdoring Adult | 0 | | 5 | | | 2.00 (0.97,3.03) | | -0.21 | | 7.07 | | -0.43 | | 1.000 |
|  | 1 | | 7 | | | 1.79 (0.91,2.66) | |  |  |  |  |  |  |  |
| !Gariep Age 6 | 0 | | 3 | | | 3.50 (2.17,4.84) | | 1.13 | | 4.99 | | 0.95 | | 0.945 |
|  | 1 | | 4 | | | 4.62 (3.47,5.78) | |  |  |  |  |  |  |  |
| !Gariep Adult | 0 | | 2 | | | 2.00 (0.36,3.64) | | 1.38 | | 3.00 | | 1.72 | | 0.650 |
|  | 1 | | 4 | | | 3.38 (2.22,4.53) | |  |  |  |  |  |  |  |
| **Supplementary Table 12: Below Umbilicus SFT and Lactation Status** | | | | | |  | |  | |  | |  | |  |
| **a) Type-II ANOVA for Lactation Status and SFT Below Umbilicus** | | | | | |  | |  | |  | |  | |  |
| SFT BU ~ Lactating*Age*Location |  | |  | | |  | |  | |  | |  | |  |
|  | **F** (16,28) | | **df** | | | **p value** | |  | |  | |  | |  |
| Lactating | 0.62 | | 1 | | | 0.437 | |  | |  | |  | |  |
| Age Category | 2.01 | | 1 | | | 0.167 | |  | |  | |  | |  |
| **Location** | **4.03** | | **1** | | | **0.055** | | * | |  | |  | |  |
| Lactating:Age Category | 0.34 | | 1 | | | 0.565 | |  | |  | |  | |  |
| Location:Age Category | 1.15 | | 1 | | | 0.293 | |  | |  | |  | |  |
| Lactating:Location | 0.34 | | 1 | | | 0.565 | |  | |  | |  | |  |
| Lactating:Location:Age Category | - | | - | | | - | |  | |  | |  | |  |
| **b) Games-Howell Post Hoc Comparisons** |  | |  | | |  | |  | |  | |  | |  |
|  | **Lactating** | | **n** | | | **Marginal Means** | | **Estimate** | | **df** | | **t** | | **p value** |
| Soetdoring Age 6 | 0 | | 10 | | | 2.05 (1.57,2.53) | | - | | - | | - | | - |
|  | 1 | | 0 | | | - | |  |  |  |  |  |  |  |
| Soetdoring Adult | 0 | | 5 | | | 1.90 (1.22,2.58) | | -2.47 | | 5.57 | | -0.66 | | 1.000 |
|  | 1 | | 7 | | | 1.57 (1.00,2.15) | |  |  |  |  |  |  |  |
| !Gariep Age 6 | 0 | | 3 | | | 3.00 (2.12,3.88) | | -0.38 | | 3.61 | | -0.54 | | 1.000 |
|  | 1 | | 4 | | | 2.62 (1.86,3.39) | |  |  |  |  |  |  |  |
| !Gariep Adult | 0 | | 2 | | | 2.00 (0.92,3.08) | | 0.13 | | 3.00 | | 0.40 | | 0.999 |
|  | 1 | | 4 | | | 2.12 (1.36,2.89) | |  |  |  |  |  |  |  |
| **Supplementary Table 13: Subscapular SFT and Lactation Status** | | | | | |  | |  | |  | |  | |  |
| **a) Type-II ANOVA for Lactation Status and SFT Subscapular** | | |  | | |  | |  | |  | |  | |  |
| SFT Subscap ~ Lactating*Age*Location |  | |  | | |  | |  | |  | |  | |  |
|  | **F** (5,28) | | **df** | | | **p value** | |  | |  | |  | |  |
| Lactating | 0.63 | | 1 | | | 0.435 | |  | |  | |  | |  |
| Age Category | 0.01 | | 1 | | | 0.905 | |  | |  | |  | |  |
| **Location** | **31.19** | | **1** | | | **< 0.001** | | *** | |  | |  | |  |
| Lactating:Age Category | 0.11 | | 1 | | | 0.742 | |  | |  | |  | |  |
| Location:Age Category | 0.33 | | 1 | | | 0.569 | |  | |  | |  | |  |
| Lactating:Location | 0.50 | | 1 | | | 0.485 | |  | |  | |  | |  |
| Lactating:Location:Age Category | - | | - | | | - | |  | |  | |  | |  |
| **b) Games-Howell Post Hoc Comparisons** |  | |  | | |  | |  | |  | |  | |  |
|  | **Lactating** | | **n** | | | **Marginal Means** | | **Estimate** | | **df** | | **t** | | **p value** |
| Soetdoring Age 6 | 0 | | 10 | | | 1.90 (1.62,2.18) | | - | | - | | - | | - |
|  | 1 | | 0 | | | - | |  |  |  |  |  |  |  |
| Soetdoring Adult | 0 | | 5 | | | 2.00 (1.60,2.40) | | 0.07 | | 6.00 | | 0.42 | | 1.000 |
|  | 1 | | 7 | | | 2.07 (1.74,2.41) | |  |  |  |  |  |  |  |
| !Gariep Age 6 | 0 | | 3 | | | 3.17 (2.65,3.68) | | -0.42 | | 4.35 | | 1.15 | | 0.890 |
|  | 1 | | 4 | | | 2.75 (2.31,3.19) | |  |  |  |  |  |  |  |
| !Gariep Adult | 0 | | 2 | | | 3.00 (2.37,3.63) | | -0.25 | | 3.00 | | 1.73 | | 0.640 |
|  | 1 | | 4 | | | 2.75 (2.31,3.19) | |  |  |  |  |  |  |  |
| **Supplementary Table 14: Mid-Biceps SFT and Lactation Status** | | | | | |  | |  | |  | |  | |  |
| **a) Type-II ANOVA for Lactation Status and SFT Mid-Biceps** | | |  | | |  | |  | |  | |  | |  |
| SFT MB ~ Lactating*Age*Location |  | |  | | |  | |  | |  | |  | |  |
|  | **F** (6,27) | | **df** | | | **p value** | |  | |  | |  | |  |
| Lactating | 0.01 | | 1 | | | 0.908 | |  | |  | |  | |  |
| Age Category | 0.46 | | 1 | | | 0.503 | |  | |  | |  | |  |
| Location | 2.73 | | 1 | | | 0.110 | |  | |  | |  | |  |
| Lactating:Age Category | 2.96 | | 1 | | | 0.096 | |  | |  | |  | |  |
| Location:Age Category | 1.11 | | 1 | | | 0.302 | |  | |  | |  | |  |
| Lactating:Location | 1.20 | | 1 | | | 0.282 | |  | |  | |  | |  |
| Lactating:Location:Age Category | - | | - | | | - | |  | |  | |  | |  |
| **b) Games-Howell Post Hoc Comparisons** |  | |  | | |  | |  | |  | |  | |  |
|  | **Lactating** | | **n** | | | **Marginal Means** | | **Estimate** | | **df** | | **t** | | **p value** |
| Soetdoring Age 6 | 0 | | 10 | | | 1.75 (1.45,2.05) | | - | | - | | - | | - |
|  | 1 | | 0 | | | - | |  |  |  |  |  |  |  |
| Soetdoring Adult | 0 | | 5 | | | 1.60 (1.18,2.02) | | -0.03 | | 9.84 | | 0.10 | | 1.000 |
|  | 1 | | 7 | | | 1.57 (1.21,1.93) | |  |  |  |  |  |  |  |
| !Gariep Age 6 | 0 | | 3 | | | 2.17 (1.62,2.71) | | -0.42 | | 2.43 | | 0.90 | | 0.950 |
|  | 1 | | 4 | | | 1.75 (1.28,2.22) | |  |  |  |  |  |  |  |
| !Gariep Adult | 0 | | 2 | | | 1.50 (0.83,2.17) | | -0.50 | | NA | | NA | | NA |
|  | 1 | | 4 | | | 2.00 (1.53,2.47) | |  |  |  |  |  |  |  |
| **Supplementary Table 15: Suprailiac SFT and Lactation Status** | | | | | |  | |  | |  | |  | |  |
| **a) Type-II ANOVA for Lactation Status and SFT Suprailiac** | | |  | | |  | |  | |  | |  | |  |
| SFT Suprailiac ~ Lactating*Age*Location |  | |  | | |  | |  | |  | |  | |  |
|  | **F** (6,27) | | **df** | | | **p value** | |  | |  | |  | |  |
| Lactating | 1.50 | | 1 | | | 0.231 | |  | |  | |  | |  |
| Age Category | 0.45 | | 1 | | | 0.508 | |  | |  | |  | |  |
| **Location** | **14.38** | | **1** | | | **< 0.001** | | *** | |  | |  | |  |
| Lactating:Age Category | 2.53 | | 1 | | | 0.123 | |  | |  | |  | |  |
| Location:Age Category | 1.91 | | 1 | | | 0.178 | |  | |  | |  | |  |
| **Lactating:Location** | **4.39** | | **1** | | | **0.045** | | * | |  | |  | |  |
| Lactating:Location:Age Category | - | | - | | | - | |  | |  | |  | |  |
| **b) Games-Howell Post Hoc Comparisons** |  | |  | | |  | |  | |  | |  | |  |
|  | **Lactating** | | **n** | | | **Marginal Means** | | **Estimate** | | **df** | | **t** | | **p value** |
| Soetdoring Age 6 | 0 | | 10 | | | 1.80 (1.49,2.11) | | - | | - | | - | | - |
|  | 1 | | 0 | | | - | |  |  |  |  |  |  |  |
| Soetdoring Adult | 0 | | 5 | | | 1.60 (1.16,2.04) | | 0.04 | | 7.93 | | 0.14 | | 1.000 |
|  | 1 | | 7 | | | 1.64 (1.27,2.01) | |  |  |  |  |  |  |  |
| !Gariep Age 6 | 0 | | 3 | | | 2.50 (1.94,3.06) | | -0.13 | | 4.33 | | 0.33 | | 1.000 |
|  | 1 | | 4 | | | 2.38 (1.89,2.86) | |  |  |  |  |  |  |  |
| !Gariep Adult | 0 | | 2 | | | 3.00 (2.31,3.69) | | -1.00 | | NA | | NA | | NA |
|  | 1 | | 4 | | | 2.00 (1.51,2.49) | |  |  |  |  |  |  |  |
